# DZNep-induced single point mutation (M236I) in poxviral 2’-O-methyltransferase enhances mRNA stability and translation efficiency

**DOI:** 10.1101/2025.03.10.642379

**Authors:** Assim Verma, Ram Kumar, Himanshu Kamboj, Garvit Kumar, Virendra Kumar Meena, Vikas Sharma, Tarun K. Bhattacharya, Bhupendra N. Tripathi, Shalini Sharma, Naveen Kumar

## Abstract

Poxviruses encode two highly conserved S-adenosylmethionine (SAM)-dependent methyltransferases for mRNA capping. While D1L facilitates translation via cap-0 formation, the role of VP39-mediated cap-1 modification has been canonically restricted to shield viral mRNA from cytoplasmic pattern-recognition receptors (PRRs). Here, we uncover a novel function of VP39 by demonstrating that prolonged exposure to the SAM cycle inhibitor DZNep selected a virus escape mutant harbouring a point mutation (M236I) in VP39. VP39_mut bypasses S-adenosylhomocysteine (SAH)-mediated feedback inhibition, enhancing 2’-O-methylation of viral transcripts and promoting their association with active translation machinery. Consequently, VP39_mut significantly improves viral protein translation, replicative fitness, and yields across species. Notably, VP39_mut-capped *in vitro* transcribed (IVT) mRNA exhibited ∼2.7-fold higher protein expression compared to VP39_wt, the standard tool for IVT applications. These findings reveal a unique resistance mechanism and provide critical insights into poxvirus mRNA biology while underscoring the potential of optimized VP39 for advancing mRNA-based therapeutics and vaccine production.

## Introduction

Poxviruses represent some of the most sophisticated viral systems, with genomes spanning 130–300 kilobase pairs and encoding approximately 200 genes^1^. This extensive genomic repertoire enables autonomous cytoplasmic replication through self-encoded machinery that includes proofreading mechanisms paralleling those of eukaryotic systems^2^. Such remarkable fidelity has led to the hypothesis that eukaryotic α-polymerase evolved from a poxvirus-like ancestor^3^.

To ensure efficient protein translation, poxviruses encode distinct S-adenosylmethionine (SAM)-dependent methyltransferases (MTases) for mRNA capping. The D1L enzyme catalyzes N7 methylation of the 5’ terminal guanosine (cap-0), a modification that stabilizes viral mRNA and facilitates nuclear export and translation initiation via interactions with host cap-binding proteins ^4,5^. Subsequently, VP39 mediates 2’-O methylation (cap-1), which serves as an immune evasion strategy by shielding viral mRNA from cytoplasmic pattern recognition receptors (PRRs) such as MDA5 and RIG-I^6,7^.

The emergence of drug-resistant viral variants has necessitated a paradigm shift in antiviral drug discovery, favouring host-directed agents (HDAs) over direct-acting antivirals (DAAs). HDAs target host factors essential for viral replication, offering broad-spectrum activity and higher genetic barriers to resistance^8–11^. Several HDAs have already been successfully repurposed or are under clinical evaluation^8,9,12^. In line with this strategy, our screening of a library of potential HDAs identified 3-Deazaneplanocin A (DZNep) as a potent antiviral agent against multiple poxviruses. DZNep targets S-adenosylhomocysteine hydrolase (ACHY), a key enzyme in the SAM cycle, disrupting SAM generation while accumulating S-adenosylhomocysteine (SAH), a potent inhibitor of SAM-dependent MTases **(Extended Data Fig. 1)**^13^.

However, prolonged DZNep exposure unexpectedly selected for a resistant virus harbouring a mutation (M236I) in VP39 (VP39_mut) which besides conferring resistance, uniquely enhanced replicative fitness, contrasting with typical resistance mutations that often incur fitness costs^14–18^. This finding uncovers previously unknown aspects of poxvirus translation mechanisms and highlights an expanded role for VP39 in mRNA biology^5,19,20^. Notably, despite encoding a cap-0 modifier (D1L) presumed to facilitate translation^5^, this enzyme remained unaltered in the DZNep-resistant mutant. Yet the viral mRNA exhibited markedly enhanced translational competency compared to both wild-type and control-passaged variants. More importantly, the emergence of this phenotype occurred in interferon-deficient Vero cells^21^, suggesting that VP39 possesses previously uncharacterized functions beyond immune evasion^7,22^.

Our findings revealed that the VP39_mut mutation not only conferred resistance to DZNep but also significantly enhanced viral fitness across evolutionarily distant species, including SARS-CoV-2. Mechanistically, this mutation circumvents SAH-mediated negative feedback inhibition, resulting in increased catalytic efficiency of VP39 and enhanced 2’-O methylation of viral transcripts. These modifications improved transcript stability and facilitated preferential recruitment to active translation machinery, leading to superior translational efficiency. Beyond its virological implications, VP39_mut-capped in vitro transcribed (IVT) mRNA exhibited markedly higher and sustained protein expression (∼2.7 fold over 10 days) compared to wild-type VP39, a standard tool for IVT applications^23–28^.

While current strategies to enhance mRNA drug efficacy focus on chemical or structural modifications, these approaches often necessitate additional manufacturing steps and yield only modest improvements in translational efficiency^29–31^. In contrast, the VP39_mut offers a facile and cost-effective alternative that simplifies the production process while reducing associated costs. By enabling sustained and enhanced protein production from IVT mRNAs without additional structural modifications, this approach addresses key challenges such as dose-dependent cytotoxicity and manufacturing scalability^32,33^.

By uncovering this novel adaptive mechanism driven by selective pressure from DZNep treatment, our study highlights both the evolutionary plasticity of poxviruses and the potential utility of optimized VP39 for biotechnological innovation.

## Results

### DZNep inhibits replication of poxviruses

To identify novel anti-poxviral agents, we screened a library of host-directed agents (HDAs) and discovered that DZNep at a non-cytotoxic concentration of 1.25 µM **(Extended Data Fig. 2a)** inhibits the replication of multiple poxviruses **(Fig 1, a-c)**. Preincubation of Buffalopox virus (BPXV) with DZNep at varying concentrations did not affect virion infectivity **(Extended Data Fig. 2b)**, indicating that the inhibitor does not compromise the viability of extracellular virions. This suggests that the antiviral activity of DZNep is mediated through targeting intracellular stages of the virus replication cycle.

**Figure 1.**
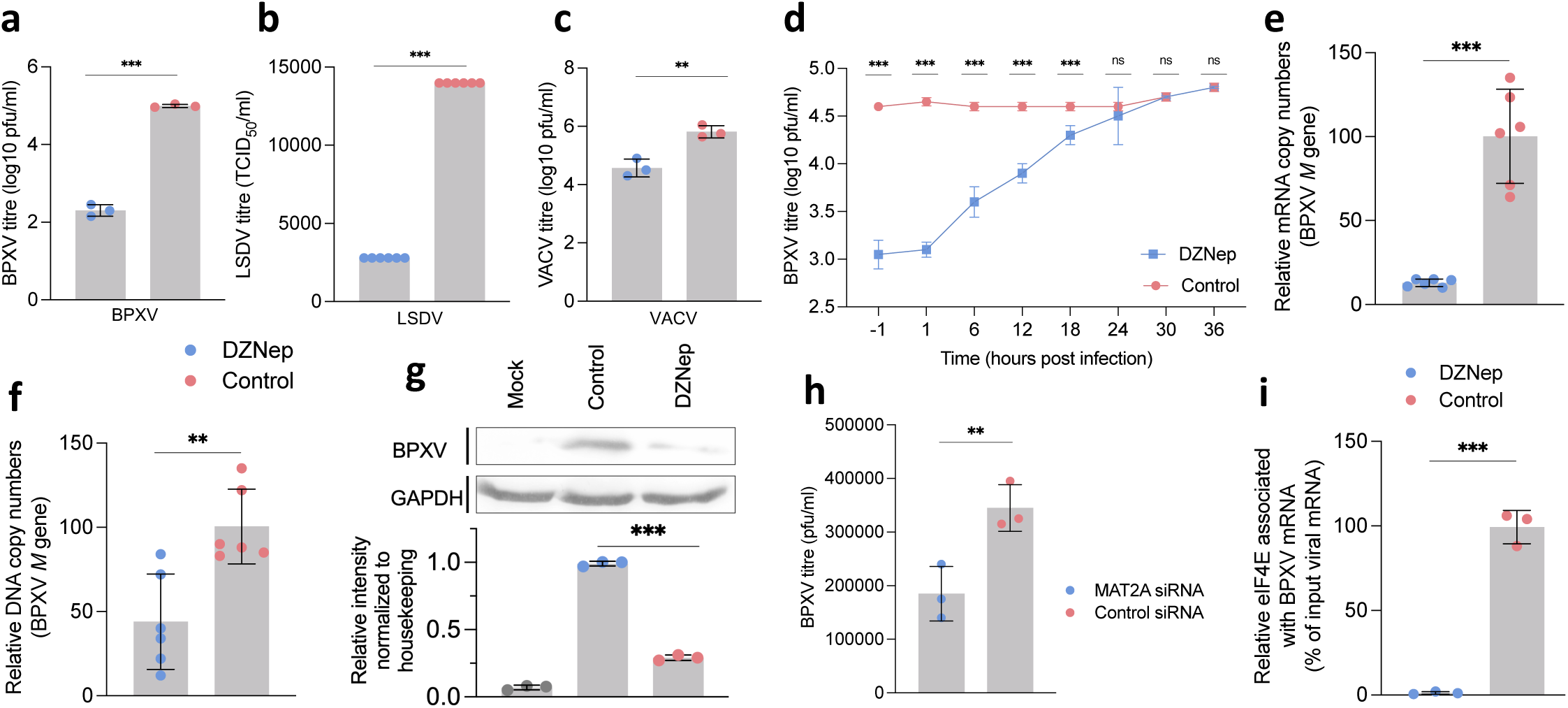
DZNep treatment suppresses poxvirus replication. a-c. *In vitro* anti-poxvirus efficacy: Vero cells were infected with the indicated viruses at an MOI of 0.1 and treated with 5 µM DZNep or DMSO in triplicates. Infectious virus particles in the culture supernatants were quantified at 72 hpi by plaque assay for BPXV and VACV, and by the TCID_50_ method for LSDV. **d. Time-of-addition assay:** Confluent monolayers of Vero cells were infected with BPXV at an MOI of 5, washed with PBS, and treated with hesperetin or DMSO at-0.5 hpi, 1 hpi, 6 hpi, 12 hpi, 18 hpi, 24 hpi, 30 hpi, and 36 hpi. Supernatants were collected at 48 hpi, and infectious virions were quantified by plaque assay. **e, f. Viral mRNA/genome synthesis:** Vero cells were infected with BPXV at an MOI of 5, washed with PBS, and treated with DZNep or DMSO at 4 hpi. RNA and DNA were isolated from cells harvested at 12 hpi and 24 hpi, respectively. For mRNA quantification, cDNA was synthesized from RNA using oligo(dT) primers. BPXV *M* gene expression was quantified by qRT-PCR, normalized to β-actin (housekeeping control), and analyzed using the ΔΔCt method. **g. Viral protein synthesis:** Vero cells were infected with BPXV and treated with DZNep or vehicle control at 4 hpi. At 24 hpi, cell lysates were prepared in RIPA buffer and analyzed by Western blot using anti-BPXV hyper-immune serum (upper panel) or GAPDH antibody (lower panel). Relative band intensities were quantified using ImageJ software and normalized to GAPDH as a loading control (bottom). **h. siRNA knockdown.** Vero cells were transfected in triplicates with MAT2A or negative control siRNAs before infection with BPXV at an MOI of 1. Virus yields in cell culture supernatants at 48 hpi were quantified by plaque assay. **i. RNA IP assay.** Vero cells were infected with BPXV at an MOI of 5 and treated with DZNep or DMSO at 4 hpi. At 16 hpi, cell lysates were prepared for RNA-IP as described in Materials and Methods. Lysates were incubated with α-peIF4E (reactive antibody), α-ERK (non-reactive antibody), or IP buffer alone (beads control), followed by Protein A Sepharose slurry incubation. After washing and cross-link reversal, cDNA was from immunoprecipitated RNA and BPXV RNA (M gene) was quantified by qRT-PCR and normalized to input controls. All values represent means ± SD from at least three independent experiments. Statistical significance was determined using Student’s t-test (**P < 0.01; ***P < 0.001).

Given the BPXV life cycle of approximately 30 hours^34^, we conducted a time-of-addition assay to pinpoint the specific stage(s) of the BPXV life cycle affected by DZNep. Both pre-treatment and treatment at 1 h post-infection (hpi) resulted in comparable inhibition of BPXV replication, suggesting that DZNep does not interfere with early stages of the viral life cycle (**Fig. 1d).** Similarly, treatment at 24 hpi had no inhibitory effect, indicating that late stages such as virion assembly and egress are unaffected. However, significant reductions in viral yield were observed when DZNep was added at 6, 12, or 18 hpi, implicating middle stages—such as genome synthesis, transcription, and translation—as the likely targets of DZNep. These findings were further corroborated by virus step-specific assays, which revealed that DZNep does not affect virus attachment, entry, or egress **(Extended Data Fig. 2c-2e).**

To evaluate the impact of DZNep on these middle stages, virus infected cells were treated with DZNep at 4 hpi (post-completion of early stages), and cell lysates were analyzed at 12 hpi. Compared to the vehicle control, cells treated with DZNep exhibited significantly reduced levels of viral mRNA **(Fig. 1e)** and DNA **(Fig. 1f)**, suggesting that DZNep targets BPXV mRNA and genome synthesis. Consistent with the correlation between reduced mRNA levels and protein expression, we also observed a decrease in the levels of viral proteins in inhibitor-treated cells compared to vehicle control-treated cells (**Fig. 1g - upper panel)**. Importantly, the levels of the housekeeping control protein GAPDH remained consistent across both groups **(Fig. 1g - lower panel)**. These findings suggested that DZNep impairs BPXV replication by targeting transcription and translation of the viral genome within the host cells.

### DZNep Disrupts Viral mRNA Cap Methylation

Unlike cap-independent translation employed by smaller viruses, such as picornaviruses, which rely on Internal Ribosome Entry Sites (IRES) within the 5’-UTR of mRNA for direct ribosome recruitment^35^, cap-dependent translation in eukaryotes and large viruses (e.g., poxviruses and coronaviruses) requires a series of tightly regulated molecular events^36^. This process is initiated by N7-methylation of the 5’-terminal guanosine (m7G) of mRNA, a critical modification mediated by the D1L MTase in poxviruses^5^. This modification facilitates interaction with eukaryotic translation initiation factor 4E (eIF4E), which subsequently drives the assembly of the eIF4F complex on the mRNA—a key step for ribosome recruitment and translation initiation^36,37^.

DZNep inhibits cellular S-adenosylhomocysteine hydrolase (ACHY), leading to restricted SAM regeneration and the accumulation of SAH. This dual effect impairs MTase activity by both depleting SAM and inducing SAH-mediated negative feedback inhibition^13^ **(Extended Data Fig. 1)**. To assess whether SAM depletion affects BPXV replication, we targeted methionine adenosyltransferase (MAT), a key enzyme in SAM biosynthesis. MAT exists in two isoforms MAT1A and MAT2A, however MAT2A has been previously shown to compensate for MAT1A loss^38^ and its inhibition lead to significant SAM depletion and metabolic disturbances^39^. As shown in **Fig. 1h**, siRNA-mediated MAT2A knockdown, significantly suppressed BPXV replication, underscoring the critical role of methylation machinery in viral propagation.

Next, RNA immunoprecipitation using α-peIF4E (phospho-eIF4E) revealed that DZNep treatment significantly reduced viral mRNA association with eIF4E compared to DMSO-treated controls **(Fig. 1i).** To verify whether this reduction is not due to the decreased p-eIF4E levels in DZNep-treated cells, we performed cell-free interaction assays between viral mRNA and p-eIF4E. Using α-peIF4E-purified peIF4E from mock-infected cells, we assessed *in vitro* interaction with normalized quantities of viral RNA isolated from DMSO-and DZNep-treated cells (quantified by qRT-PCR). The amount of RNA immunoprecipitated by the α-peIF4E complex was significantly reduced in the DZNep-treated group compared to DMSO controls **(Extended Data Fig. 2f)**. These findings confirm that DZNep-mediated inhibition of methylation disrupts 5’ cap formation on BPXV mRNA, thereby preventing its interaction with eIF4E and impairing subsequent viral protein translation.

### DZNep pressure selects virus escape mutant

To evaluate the potential for drug resistance development, BPXV was serially passaged 50 times in the presence of either DZNep or vehicle control (DMSO) **(Fig 2a – left panel)**. While both wild-type BPXV (wtBPXV) and control-passaged BPXV (dsBPXV) remained susceptible to DZNep treatment, passaged BPXV under drug pressure (drBPXV) exhibited complete resistance to DZNep inhibition **(Fig 2a – right panel).**

**Figure 2.**
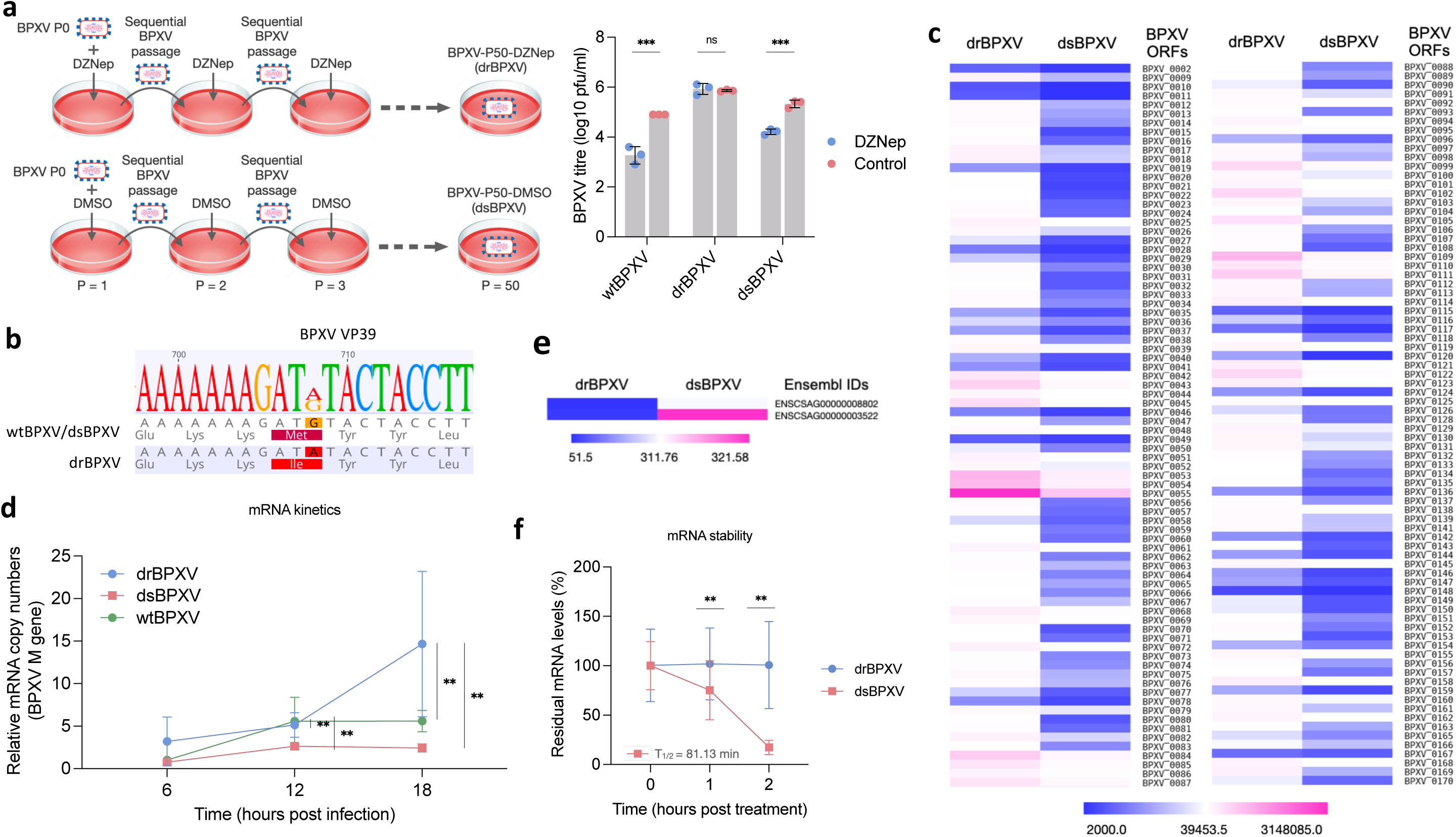
DZNep pressure selects virus escape mutant a. Selection of DZNep-resistant BPXV mutants: Vero cells were infected with BPXV at an MOI of 0.1 and treated with either DZNep (1.25 µM) or DMSO. Progeny virus particles in the supernatant were harvested at 48–72 hpi or when ∼75% of cells exhibited cytopathic effects (CPE). Fifty sequential passages were performed. Subsequently, Vero cells, in triplicate, were infected with P0 (wtBPXV), P50-DZNep (drBPXV), or P50-Control (dsBPXV) passaged viruses at an MOI of 0.1 in the presence of either DZNep (5 µM) or DMSO. Progeny virus particles released at 72 hpi were quantified by plaque assay. **b. Mutational analysis of drBPXV variant:** Whole genome sequences of wtBPXV, drBPXV, and dsBPXV were analyzed using Geneious Prime software to identify nucleotide sequence variations. **c. Viral transcript abundance:** The relative abundance of BPXV ORFs was retrieved from RNA-seq data to compare transcript levels between variants. **d. mRNA kinetics:** Vero cells, in triplicate, were infected with the indicated BPXV variant at an MOI of 5 for 1 h, followed by five PBS washes and supplementation with fresh DMEM. Cells were harvested at specified time points, RNA was isolated, and cDNA was synthesized using oligo(dT) primers. BPXV M gene expression was quantified by qRT-PCR, normalized to β-actin (housekeeping control), and relative fold-change was calculated using the ΔΔCt method. **e**. **KEGG pathway analysis:** Differentially expressed host genes involved in mRNA degradation pathways were identified from RNA-seq data using KEGG pathway filtering. **f. Measurement of mRNA stability:** Vero cells, in triplicate, were infected with BPXV at an MOI of 5. At 12 hpi, cells were treated with DZNep or DMSO in the presence of Actinomycin D (5 µg/ml). RNA was isolated at specified time points post-treatment and subjected to cDNA synthesis. BPXV M gene expression was quantified by qRT-PCR, and relative viral mRNA levels at 0 h, 1 h, and 2 h post-treatment are shown. All values represent means ± SD from at least three independent experiments. Statistical significance was determined using Student’s *t*-test (*ns* = non-significant; \*\**P* < 0.01; \*\*\**P* < 0.001).

Whole-genome sequencing revealed a point mutation in VP39 (encoding the viral 2’O-methyltransferase) in drBPXV (GenBank: PQ361283) but not in wtBPXV (GenBank: MW883892) or dsBPXV (GenBank: PQ361284) genomes **(Fig. 2b).** This mutation resulted in an amino acid substitution (M236I; protein_id=“<u>QCY54144.1</u>”), potentially conferring resistance to SAH-mediated feedback inhibition. Additional mutations were identified in both drBPXV and dsBPXV (Table 1, Table 2), likely resulting from extended cell culture adaptation.

**Table 1.**
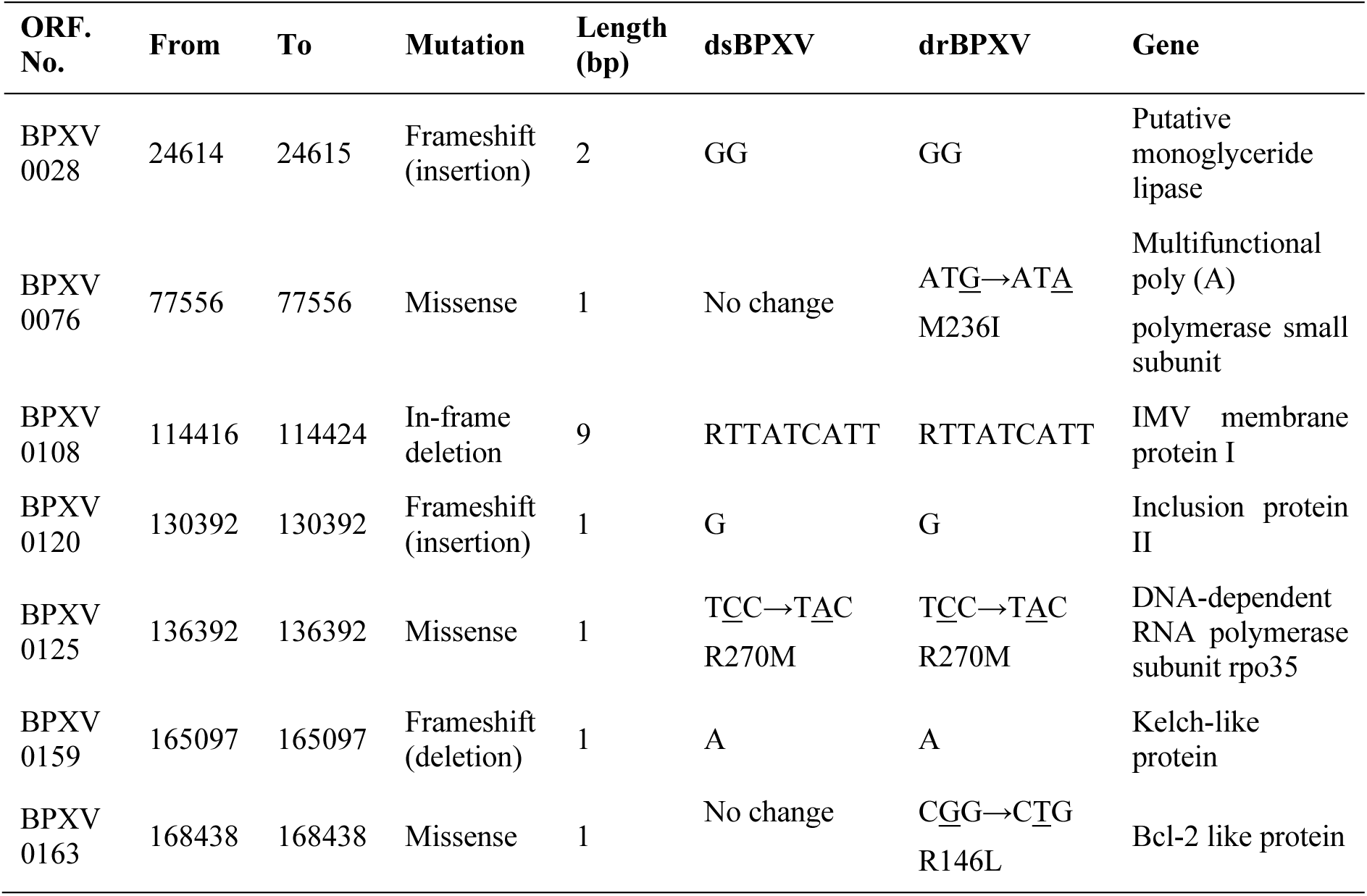
List of mutations in coding region (mapped to BPXV-WT, GenBank: MW883892)

**Table 2.** List of mutations in non-coding and ITR regions (mapped to BPXV-WT, GenBank: MW883892) coding coding coding coding coding coding coding coding coding coding coding coding coding coding.

| Region | From | To | Mutation | Length<br>(bp) | dsBPXV | drBPXV |
| --- | --- | --- | --- | --- | --- | --- |
| ITR | 6537 | 6537 | Deletion | 1 | T | T |
| Non-coding | 31067 | 31070 | Deletion | 4 | ATAT | ATAT |
| Non-coding | 33321 | 33321 | Insertion | 1 | T | No change |
| Non-coding | 137953 | 137953 | Deletion | 1 | T | T |
| Non-coding | 142094 | 142094 | Deletion | 1 | G | G |
| Non-coding | 142254 | 142259 | Deletion | 6 | AAGAWY | AAGAWY |
| Non-coding | 146279 | 146281 | Insertion | 3 | ATG | ATG |
| Non-coding | 148634 | 148634 | Deletion | 1 | T | T |
| Non-coding | 149378 | 149378 | Deletion | 1 | T | T |
| Non-coding | 149405 | 149405 | Deletion | 1 | A | A |
| Non-coding | 165686 | 165697 | Deletion | 12 | TACAGATACAGA | TACAGATACAGA |
| Non-coding | 166831 | 166831 | Deletion | 1 | T | T |
| Non-coding | 167928 | 167928 | Insertion | 1 | T | T |
| Non-coding | 169117 | 169118 | Insertion | 2 | AT | AT |
| Non-coding | 173603 | 173605 | Insertion | 3 | ATT | ATT |

### Transcriptomic signatures of resistant mutant

To elucidate the resistance mechanism, we conducted RNA-seq analysis of Vero cells infected with either drBPXV or dsBPXV **(Extended Data Fig. 3)**. Mapping statistics revealed a substantial enrichment in viral mRNA reads in drBPXV (60.95%) compared to dsBPXV (18.23%), with a corresponding decrease in host genome (*C. sabaeus*) mapping (72.83% vs 90.65%, respectively). This observation prompted us to examine the abundance of BPXV open reading frames (ORFs) in both infection groups. As illustrated in the heatmap in **Fig. 2c**, the majority of drBPXV ORFs were significantly more abundant than those in dsBPXV. To confirm this we conducted mRNA kinetics study which suggested drBPXV transcripts were significantly more abundant at 12 and 18 hpi, as compared to both wtBPXV and dsBPXV **(Fig. 2d).**

Given the role of 2’O-Me modification in preventing mRNA degradation^40,41^, we analyzed host RNA decay pathways in the RNA-seq data. Key components of the exosome complex responsible for mRNA degradation, Rrp41 and Rrp46 (Ensembl IDs: ENSCAG00000008802, ENSCAG00000003522)^42^, were significantly downregulated in drBPXV-infected cells **(Fig. 2e)**. This suggests that drBPXV transcripts evade activation of the host mRNA surveillance system. Consistent with this observation, RNA polymerase inhibitor, actinomycin D-mediated mRNA stability assay showed prolonged stability of drBPXV transcripts, whereas dsBPXV transcripts exhibited progressive decay with a half-life of 81.13 minutes **(Fig. 2f)**.

Furthermore, KEGG pathway analysis revealed significant alterations in cellular pathways related to metabolism, ribosomal biogenesis, and metabolite biosynthesis in drBPXV-infected cells compared to dsBPXV-infected cells **(Extended Data Fig. 4)**. These findings collectively suggest that drBPXV transcripts are remarkably stable which may lead to enhanced hijacking of host machinery.

### drBPXV acquired enhanced replicative fitness

In addition to modest titre enhancement in both drBPXV and dsBPXV compared to wtBPXV, potentially attributable to cell culture adaptation, drBPXV exhibited significantly higher titers than dsBPXV irrespective of DZNep treatment **(Fig. 2a, right panel)**. Similar pattern was also observed in western blot analysis, confirming the elevated protein production in drBPXV-infected cells compared to dsBPXV-infected cells **(Fig. 3a)**. Moreover, one-step growth curves also demonstrated accelerated replication kinetics of drBPXV compared to wild-type and dsBPXV variants **(Fig. 3b)**. Remarkably, drBPXV formed larger plaques **(Fig. 3c)** and induced more pronounced cytopathic effects (CPE) irrespective of DZNep treatment **(Extended Data Fig. 5)**, suggestive of the acquisition of enhanced replicative fitness.

**Figure 3.**
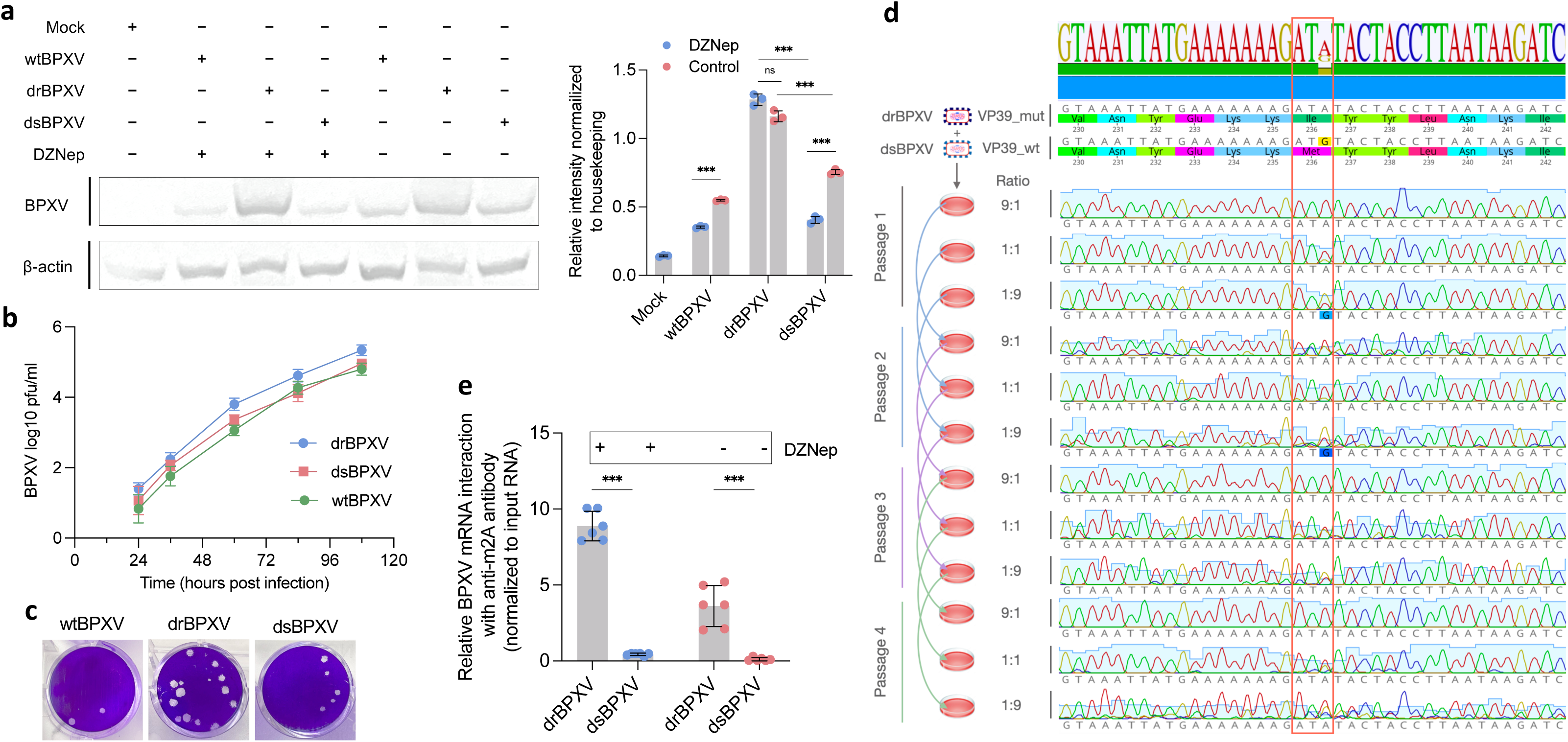
drBPXV exhibits enhanced replicative fitness a. Viral protein synthesis: *Left panel* - Vero cells were infected with the indicated BPXV variants or mock-infected and treated with DZNep or vehicle control at 3 hpi. At 24 hpi, cell lysates were prepared in RIPA buffer and analyzed by western blot using anti-BPXV hyper-immune serum (upper panel) and GAPDH antibody (lower panel) as a housekeeping control. *Right panel* - Relative band intensities, normalized to GAPDH (loading control), were quantified using ImageJ software. **b. One-step growth curves**. Confluent monolayers of Vero cells, in triplicates, were infected with BPXV at an MOI of 0.1 for 1 h, followed by PBS washing and addition of fresh medium. Infectious progeny virus particles in the supernatants were quantified at the indicated time points by plaque assay. Results are representative of three independent experiments. **c. Plaque morphology:** Representative images of plaque morphology for wtBPXV, drBPXV, and dsBPXV variants. **d. Direct competition assays:** To compare the relative fitness of BPXV-P50-DZNep (drBPXV) and BPXV-P50-control (dsBPXV), both variants were mixed at ratios of 9:1, 1:1, or 1:9 and inoculated onto Vero cells at an MOI of 0.1. After observing CPE, supernatants were harvested and sequentially passaged up to passage 4 (p = 4). Each passaged variant was PCR-amplified using high-fidelity polymerase targeting the region harbouring M236I mutation in VP39, followed by Sanger sequencing. The abundance of each competitor was determined by measuring the height of the nucleotide encoding WT (nucleotide G) or mutant (nucleotide A) in sequencing chromatograms. **e. RNA IP assay:** Vero cells were infected with BPXV at an MOI of 5 and treated with DZNep or DMSO at 6 hpi. At 16 hpi, cell lysates were prepared for RNA-IP as described in Materials and Methods. Lysates were incubated with α-m2A (reactive antibody), α-ERK (nonreactive antibody), or IP buffer alone (beads control), followed by Protein A Sepharose slurry incubation. After washing and cross-link reversal, cDNA was synthesized from immunoprecipitated RNA, and BPXV RNA (M gene) was quantified by qRT-PCR and normalized to input RNA. Values are means ± SD and representative of the result of at least 3 independent experiments. Pair-wise statistical comparisons were performed using the Student’s t test (ns = non-significant, *** = P<0.001).

To directly assess competitive fitness, we conducted a direct-competition assay between drBPXV and dsBPXV. Analysis of the chromatograms of nucleotide sequencing centered around M236I mutation of VP39 **(Fig. 3d)** revealed that drBPXV outcompeted dsBPXV within the first passage at 9:1 and 1:1 ratios, and by the third passage at a 1:9 ratio, demonstrating superior replicative fitness.

### VP39_mut bypasses SAH-mediated negative feedback inhibition

The observed enhancement in transcript stability **(Fig. 2f)** and abundance **(Fig. 2c)** in drBPXV prompted us to hypothesize that VP39_mut exhibits higher catalytic efficiency. To confirm this, we performed RNA immunoprecipitation in virus infected lysates using α-m2A. As shown in **Fig. 3e**, drBPXV transcripts demonstrated significantly higher enrichment of 2’O-Me marks compared to dsBPXV **(Fig. 3e)**. Notably, DZNep treatment did not affect this enrichment in drBPXV, explaining the observed insusceptibility of drBPXV in antiviral **(Fig. 2a)** and western blot assays (**Fig. 3a)**. Given that SAH is a potent inhibitor of VP39 activity even at sub-stoichiometric concentrations relative to SAM^13^, we further investigated the structural basis of VP39_mut’s apparent resistance to feedback inhibition.

Molecular dynamics (MD) simulations revealed distinct conformational dynamics between VP39_wt and VP39_mut. While the VP39_wt-SAH complex maintained stable ligand binding throughout 200 ns **(Fig. 4a),** VP39_mut underwent conformational changes displacing SAH by 150 ns **(Fig. 4b, Supplementary movie 1)**. Binding free energy calculations corroborated this observation, with VP39_mut-SAH showing reduced affinity (ΔG:-5.3 kcal/mol) compared to VP39_wt-SAH (-7.4 kcal/mol) due to fewer stabilizing interactions **(Fig. 4c, Extended Data Fig. 6g, 6h)**. Root mean square deviation (RMSD) analysis demonstrated increased conformational flexibility in VP39_mut after SAH displacement at ∼150 ns **(Fig. 4f)**, while diffusion maps highlighted enhanced atomic fluctuations post-displacement **(Fig. 4d, 4e)**. Root mean square fluctuation (RMSF) calculations identified heightened flexibility within residues spanning positions 210–240, encompassing the mutation site, in VP39_mut compared to VP39_wt **(Fig. 4g)**.

**Figure 4.**
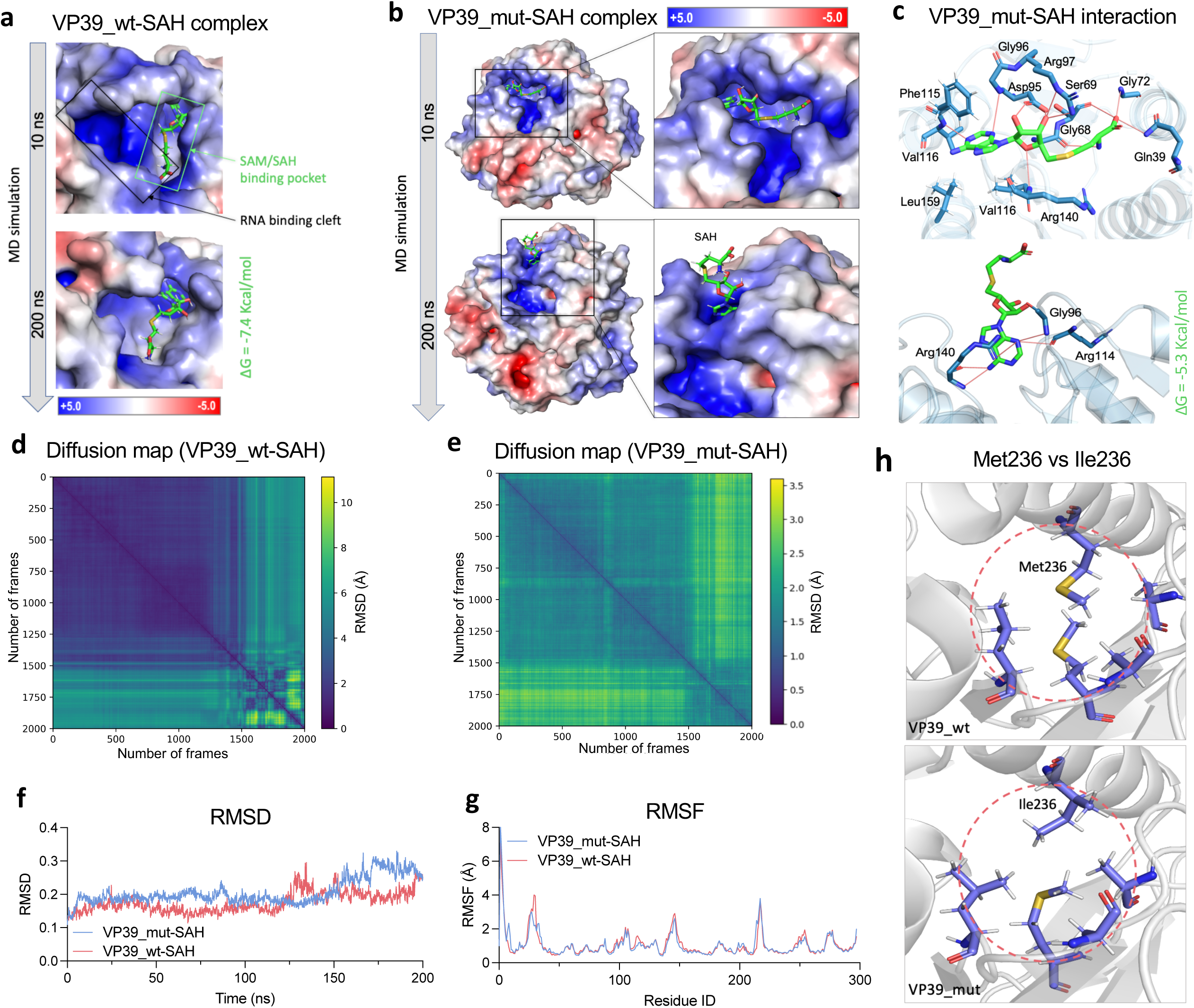
VP39_mut circumvents SAH-mediated negative feedback inhibition a, b. Molecular dynamics (MD) simulation of VP39-SAH complex: The crystal structure of the VP39-SAH complex (PDB: 1VP3) was used for 200 ns MD simulations. The surface of the VP39 protein is colored according to the electrostatic surface potential. **c. Interaction analysis:** Interacting residues of VP39_mut with SAH at 10 ns (*upper panel*) and 200 ns (*lower panel*). Hydrogen bonds are depicted as red lines. **d, e. Diffusion maps:** Diffusion maps based on distance matrices for VP39_wt and VP39_mut complexes with SAH during the simulation. **f. RMSD plot:** Root mean square deviation (RMSD) analysis of VP39_wt and VP39_mut in complex with SAH over the course of the simulation. **g. RMSF plot:** Root mean square fluctuation (RMSF) analysis of VP39_wt and VP39_mut in complex with SAH, highlighting differences in residue flexibility during the simulation. **h. Structural implications of M236I mutation:** *Upper panel* - Met236 in the VP39_wt structure; *Lower panel* - Ile236 mutation in the VP39_mut structure. Surrounding amino acid residues are shown as blue stick representations in both structures.

Simulations with SAM-bound complexes revealed enhanced structural stability in VP39_mut **(Extended Data Fig. 6a-c)** compared to the sustained fluctuations in VP39_wt **(Extended Data Fig. 6d-e),** particularly in the 210-240 alpha-helical region **(Extended Data Fig. 6f)**. The M236I substitution, replacing bulky thioether group of methionine with compact side chain of isoleucine **(Fig. 4h)**, reduced steric hindrance and enhanced VP39_mut-SAM stability (ΔG:-8.5 kcal/mol vs-7.5 kcal/mol for VP39_wt-SAH) **(Extended Data Fig. 6i, 6j)**.

These structural insights demonstrate that the M236I mutation selectively disrupts SAH binding while maintaining SAM affinity, thereby enabling enhanced catalytic activity despite elevated SAH levels.

### VP39_mut augments protein translation

The emergence of the resistance mutation in VP39 within interferon-deficient Vero cells, coupled with its enhancement of viral protein synthesis, led us to hypothesize that VP39 activity extends beyond immune evasion and potentially contributes to translational competency. Polysome profiling revealed significantly higher polysome peaks in drBPXV-infected cells compared to dsBPXV-infected cells, indicative of enhanced active translation **(Fig. 5a)**. qRT-PCR analysis of viral mRNA distribution across monosome and polysome fractions demonstrated significant enrichment of drBPXV transcripts in polysome fractions **(Fig. 5b)**. These findings, combined with RNA immunoprecipitation data **(Fig. 3e)**, suggest that the enhanced 2’-O-methylation activity of VP39_mut facilitates increased translational capacity.

**Figure 5.**
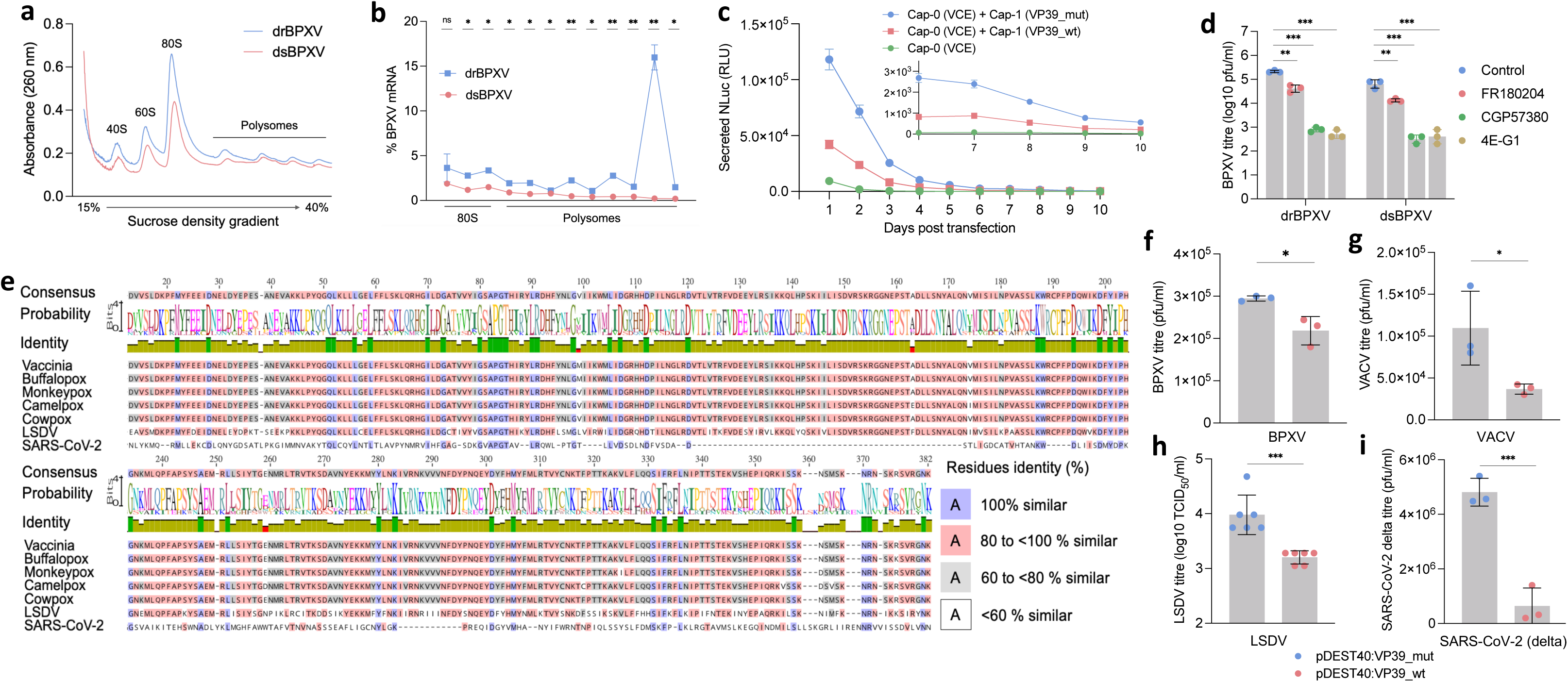
VP39_mut enhances translation a. Polysome profiling: Vero cells were infected with dsBPXV or drBPXV at an MOI of 5 for 12 h. Cell lysates were layered onto sucrose gradients at normalized OD260 and fractionated by sucrose density gradient velocity sedimentation. **b. qRT PCR of mono-and polysome fractions:** RNA was isolated from the retrieved m_0_ ono-and polysome fractions, followed by cDNA synthesis using oligo(dT) primers. Viral (M gene) and housekeeping (β-actin) transcripts were quantified by qRT-PCR. **c. Time-course IL6-Nanoluciferase secretion.** IL6-Nanoluciferase mRNA constructs, capped with vaccinia capping enzyme (cap-0) were subjected cap-1 modification either with VP39_wt or VP39_mut followed by transfection in HeLa cells in triplicates. Luminescence were measured daily for 10 days post-transfection using Spectramax i3x in end-point mode. Each day complete medium was harvested and replaced with fresh 150 µl media and 10 µl of harvested media was used for luciferase assay. **d. ERK-MNK1-eIF4E signaling is a prerequisite for drBPXV:** Vero cells were infected with drBPXV or dsBPXV at an MOI of 0.1. At 1 hpi, cells were washed with PBS, and fresh medium containing an ERK inhibitor (FR180204: 0.2 μg/ml), MNK1 inhibitor (CGP57380: 0.5 μg/ml) or eIF4E inhibitor (4EGI-1; 0.5 μg/ml) was added. The progeny virus particles released in the infected cell culture supernatant at 48 hpi were quantified by plaque assay. **e. 2’O-MTase sequence conservation: T**he 2’O-MTase sequences from poxviruses and SARS-CoV-2 were retrieved from GenBank and aligned using MUSCLE in Geneious Prime software (2024.0.7). **f-i. VP39 overexpression:** HeLa cells (for BPXV, LSDV, and VACV) and A549 cells (for SARS-CoV-2) were transfected in triplicates with pDEST40 plasmid constructs encoding VP39_WT or VP39_mut. After 48 h, cells were infected with the indicated viruses at an MOI of 5. Supernatants were collected at 42 hpi (BPXV, LSDV, VACV) or 24 hpi (SARS-CoV-2), and infectious progeny viruses were quantified using plaque assays on Vero cells. Values are means ± SD and representative of the result of at least 3 independent experiments. Pair-wise statistical comparisons were performed using the Student’s t test (ns = non-significant, * = P<0.05, ** = P<0.01, *** = P<0.001).

To evaluate whether VP39_mut could enhance translation in current state-of-the-art synthetic mRNA platform, we performed in vitro transcription experiments using IL6-Nanoluciferase mRNA capped at the cap-0 position using vaccinia capping enzyme, followed by cap-1 methylation with either VP39_wt or VP39_mut. The capped mRNAs were transfected into HeLa cells, and luminescence was measured at 24-hour interval over 10 days. Remarkably, transcripts capped by VP39_mut exhibited a ∼2.8-fold increase in luminescence at 24 hours post-transfection compared to those capped by VP39_wt **(Fig. 5c)**. Notably, this enhancement persisted through day 10, with VP39_mut-capped mRNA maintaining ∼2.6-fold higher luminescence. In contrast, cap-0 modified mRNA displayed significantly reduced luminescence (∼4.5-and ∼12.6-fold lower compared to VP39_wt-and VP39_mut-capped mRNAs, respectively). Besides demonstrating a direct translational role of VP39, these results highlight the potential utility of VP39_mut in applications requiring robust and prolonged gene expression, such as mRNA-based vaccines and gene delivery systems.

### VP39_mut mediated translation is ERK/MNK/eIF4E dependent

Poxviruses employ canonical cap-dependent translation mediated through the ERK/MNK/eIF4E signaling axis. In this pathway, extracellular signal-regulated kinase (ERK) activates MAP kinase-interacting kinases (MNKs), which phosphorylate eIF4E—a critical step for recognizing the cap-0 structure on mRNA. This recognition facilitates the assembly of the eIF4F complex (comprising eIF4E, eIF4G, and eIF4A), which recruits the 43S pre-initiation complex to the mRNA, enabling ribosome scanning and translation initiation.

To investigate whether drBPXV employs an alternative non-canonical route for translation initiation, potentially bypassing eIF4E dependency^28^, particularly given that cap recognition by eIF4E is a rate-limiting step in translation initiation^43^, we conducted antiviral assays using small-molecule inhibitors targeting key components of the ERK/MNK/eIF4E pathway: FR180204 (ERK inhibitor), CGP57380 (MNK1 inhibitor), and 4EGI-1 (eIF4E inhibitor)^44^. As shown in **Fig. 5d**, both dsBPXV and drBPXV exhibited similar susceptibility to these inhibitors, suggesting that the translational enhancement mediated by VP39_mut operates downstream of the ERK/MNK/eIF4E signalling axis. These findings align with a recent study demonstrating that chemically introduced 2’-O-methylation at the 5’-end of transcripts enhances eIF4E binding affinity and promotes the formation of a stable RNA–eIF4E–eIF4G ternary complex, thereby facilitating efficient translation initiation^30^.

### VP39_mut increase titres of diverse viruses

VP39 is highly conserved among poxviruses, with orthopoxviruses (BPXV, Mpox, VACV, and CPXV) sharing identical VP39 sequences, while LSDV, a capripoxirus, maintains 92.5% amino acid identity **(Fig. 5e, Extended Fig. 7)**. Recent structural analyses have further revealed that VP39 conservation extends to evolutionarily distant viruses, including coronaviruses^45,46^. Notably, even minor natural mutations in 2’O-methyltransferases (2’O-MTases) among closely related viruses has profound influence on catalytic activity, thereby modulating virulence and pathogenesis^45^.

To explore the cross-functional potential of VP39_mut, we conducted overexpression studies across diverse viral species. Remarkably, VP39_mut significantly enhanced viral titers of poxviruses (BPXV, VACV, LSDV) as well as SARS-CoV-2 **(Fig. 5f-i).** Intriguingly, the enhancement was even more pronounced in SARS-CoV-2 (∼7.5-fold increase), suggesting that VP39_mut exhibits broad functional compatibility across evolutionarily divergent viruses. These findings underscore the potential application of VP39_mut for improving viral yields in cell culture systems, which could be particularly valuable for vaccine production and other virological applications.

## Discussion

The presence of functionally redundant proteins within space-constrained viral genomes presents an intriguing paradox in viral biology^47^. Our study reveals how prolonged DZNep pressure drives poxviruses to optimize their replication strategy by shifting reliance from the cap-0 modifying enzyme (D1L) to the cap-1 modifying enzyme (VP39) for enhanced protein translation. This rare adaptive mechanism provides novel insights into previously unexplored aspects of poxvirus replication.

Unlike smaller eukaryotic viruses, poxviruses encode an extensive repertoire of genes within their exceptionally large genomes, enabling complex replication strategies with built-in redundancies. Although experimental and technical limitations have hindered a comprehensive understanding of these strategies, poxviruses are known to utilize both cap-dependent (cap-0) and cap-independent translation mechanisms^48,49^. The latter mechanism, uniquely observed in poxviruses^50,51^, utilizes poly(A) leader sequences in post-replicative mRNAs to selectively drive translation following decapping, which effectively suppresses host protein synthesis^52^. Although efficient, temporal restriction of this mechanism necessitates cap-dependent translation during early replication stages. This dual strategy ensures that poxviruses meet the extensive demands for protein production across different stages of their life cycle.

Our findings suggest that under DZNep pressure, drBPXV preferentially optimizes VP39 activity over D1L. This shift highlights a critical gap in our understanding of poxvirus gene expression and underscores the fundamental role of VP39 in translation. We hypothesize that this adaptation may be driven by the rate-limiting nature of eIF4E-cap interactions, as eIF4E is typically present in limited quantities^43^. Although alternative cap-binding proteins such as eIF3d have been shown to compensate for eIF4E loss through non-canonical translation initiation^53^, this mechanism requires specific mRNA features (e.g., 5’-UTR stem-loops) and operates under restricted cellular conditions^54^. Additionally, D1L MTase activity for cap-0 maturation depends on its association with the D12L subunit^55^, a two-subunit dependency that likely complicates resistance development through mutation. These host-and poxvirus-specific constraints may have driven drBPXV to optimize its translation strategy through VP39.

Although VP39 activity has traditionally been associated with cap-1 modification^56,57^, its sequence and structural conservation extend to evolutionarily distant species, such as *Trypanosoma*, where it catalyzes the methylation of up to four transcribing nucleotides (cap-4)^58,59^. However, technical limitations in mapping 2’-O-methylation of mRNA at single-nucleotide resolution^60–62^ have hindered a comprehensive understanding of VP39 capability and the biological significance of multiple cap modifications. Notably, mammalian mRNA utilizes cap-2 modifications to enhance stability and translational efficiency compared to cap-1 structures^63^. Notably, VP39 also shares homology with the human-encoded 2’O-MTase, suggesting evolutionary conservation of this mechanism across diverse taxa^64^.

These observations raise the intriguing possibility that poxviruses may exploit higher-order cap structures to optimize translation. Notably, recent studies employing chemically modified IVT mRNA with up to cap-5 structures increased affinity for eIF4E and eIF4G and demonstrated significantly enhanced protein expression^30^. Consistent with these findings, our polysome profiling data revealed a robust association between VP39_mut-modified mRNA and active translation machinery, highlighting the critical translational advantage conferred by VP39 optimization.

This translational advantage was further evident in overexpression studies, wherein VP39_mut significantly enhanced titres of poxviruses and SARS-CoV-2. Notably, the fold increase in SARS-CoV-2 titers was substantially higher compared to poxviruses. Given that SARS-CoV-2 encodes only 29 genes, in contrast to the hundreds of transcripts requiring capping in poxviruses, this pronounced effect likely reflects the high catalytic efficiency of VP39_mut under conditions of reduced substrate demand.

These findings hold significant biotechnological implications. Besides demonstrating cross-functional utility in improving viral titres of evolutionary distant viruses, our data indicate that the increased capping efficiency of VP39_mut can be leveraged to improve the efficacy of current mRNA platforms. Importantly, VP39_wt exhibits limited catalytic efficiency^65^, necessitating higher mRNA doses—a factor potentially associated with increased cytotoxicity^32^. This positions VP39_mut as a superior alternative for reducing mRNA dosage requirements and mitigating cytotoxicity in therapeutic applications.

In conclusion, these unprecedented findings offer potential to improve both traditional vaccines as well as cutting-edge mRNA-based therapeutics. By uncovering a novel viral resistance mechanism to an epidrug, this serendipitous discovery of VP39_mut opens new avenues for safer and more effective therapeutic strategies.

## Author contributions

Conceptualization, N.K., A.V. Methodology and investigation, A.V., R.K. Cloning and Biochemical assays, A.V., H.K., G.K. *in silico* work, A.V., V.K.M, V.S. Data analysis, T.K.B., B.N.T, S.S., N.K. Supervision, N.K. Writing-original draft, A.V., Writing-review and editing, A.V., N.K.

## Materials and methods

### Cells and viruses

Mycoplasma-free African green monkey kidney (Vero) and HeLa cells from the National Centre for Veterinary Type Cultures (NCVTC), Hisar were maintained in Dulbecco’s Modified Eagle’s Medium (DMEM) supplemented with antibiotics and 10% fetal calf serum. The following viruses were used: Vero cell-adapted BPXV (VTCC-AVA90), A549-adapted SARS-CoV-2 Delta variant (VTCC-AVA319), and Vero cell-adapted lumpy skin disease virus (LSDV) (VTCC-AVA288) from the NCVTC repository, and Vaccinia virus from the American Type Culture Collection (ATCC). Viral titers were determined using TCID50/ml for LSDV and plaque assays (PFU/ml) for BPXV, SARS-CoV-2, and Vaccinia virus.

### Inhibitor

3-Deazaneplanocin (DZNep) was procured from BioGems International Inc. (Westlake Village, CA, USA) and was dissolved in dimethyl sulfoxide (DMSO). FR180204, CGP57380 and 4EGI-1 were procured from Sigma (Steinheim, Germany). The subcytotoxic concentration of ERK inhibitor (FR180204), MNK1 inhibitor (CGP57380) and eIF4E inhibitor (4EGI-1) were 0.2 μg/ml, 0.5 μg/ml and 0.5 μg/ml, respectively, and have been described previously^66^.

### Antibodies

Primary antibodies included eIF4E monoclonal antibody (5D11), anti-6xHis antibody (Invitrogen, South San Francisco, CA, USA), mouse anti-GAPDH primary antibody (housekeeping control), and rabbit recombinant monoclonal 2-methyladenosine (m2A) antibody (Abcam, UK). Secondary antibodies included goat anti-mouse IgG-alkaline phosphatase and goat anti-rabbit IgG-peroxidase (Sigma-Aldrich, St. Louis, USA). BPXV-specific rabbit hyperimmune serum was obtained from NCVTC, Hisar^67^.

### Cytotoxicity and virucidal activity of DZNep

The cytotoxicity (MTT assay) and virucidal effect of DZNep were evaluated as described previously^68^.

### *In vitro* antiviral efficacy

Vero cells were infected with BPXV (MOI 0.1) in triplicate and treated with DZNep (0.56-5 µM) or DMSO (vehicle control). After 1 hour of infection, cells were washed with PBS and replenished with fresh DMEM containing DZNep or DMSO. Viral titers in culture supernatants were quantified by plaque assay at 72 hours post-infection (hpi). The effective concentration 50 (EC50) of DZNep was calculated using the Reed-Muench method^69^.

### Time-of-addition assay

Confluent monolayers of Vero cells, in triplicates, were infected with BPXV at MOI of 5 for 1 h, followed by five times washing with PBS and addition of DZNep (5 µM) or 0.05% DMSO (vehicle control) at-0.5 hpi, 1 hpi, 6 hpi, 12 hpi, 18 hpi, 24 hpi and 30 hpi. The supernatants from the virus infected cells were collected at 36 hpi and quantified by plaque assay.

### Virus step-specific assays

For attachment assay, Vero cell monolayers were pre-treated with DZNep (5 µM) or vehicle control for 1 hour, followed by BPXV infection (MOI 5) at 4°C for 1 hour. After five PBS washes to remove unbound virus, cell lysates were prepared by freeze-thaw cycling and viral titers were quantified by plaque assay.

For virus entry studies, prechilled Vero cell monolayers were infected with BPXV (MOI 5) at 4°C for 1 hour to permit attachment. Following five PBS washes, cells were incubated with DZNep or vehicle control at 37°C for 1 hour to allow virus entry. After removing extracellular virus by PBS washing, cells were maintained in inhibitor-free DMEM. Infectious virus particles in supernatants were quantified by plaque assay at 48 hpi.

To assess virus release, BPXV-infected Vero cells (MOI 5) were incubated until 36 hpi. Cells were then washed and treated with DZNep or vehicle control. Supernatants were collected at 30 minutes and 4 hours post-treatment for plaque assay quantification.

For viral protein synthesis analysis, BPXV-infected (MOI 5) or mock-infected Vero cells were treated with DZNep or vehicle control at 3 hpi. Cell lysates were prepared at 24 hours post-infection for western blot analysis of viral and cellular proteins.

To quantify viral RNA/DNA, BPXV-infected Vero cells (MOI 5) were treated with DZNep or vehicle control at 3 hours post-infection. At 12 hours post-infection, total RNA/DNA was extracted. cDNA was synthesized using oligo dT primers (Fermentas, Hanover, USA) for mRNA quantification. BPXV M gene and β-actin expression were quantified by qRT-PCR as described previously^68^.

### RNA immuno-precipitation (RNA-IP) assay

RNA-IP was performed to assess viral mRNA interaction with eIF4E and evaluate 2’-O-methylation levels in viral transcripts^70^.

Vero cells were infected with BPXV (MOI 5) and treated with DZNep or vehicle control at 3 hours post-infection. At 12 hours post-infection, protein-nucleic acid complexes were cross-linked with 1% formaldehyde for 10 minutes and quenched with 125 mM glycine. Cells were lysed in immunoprecipitation buffer (150 mM NaCl, 50 mM Tris-HCl [pH 7.5], 5 mM EDTA, 0.5% NP-40, 1% Triton X-100, and protease/phosphatase inhibitor cocktail).

Cell lysates were sonicated (Qsonica Q500) using six 15-second pulses at 40% amplitude, clarified by centrifugation (12,000 g, 10 minutes), and supplemented with 10 units of RiboLock RNase Inhibitor (Invitrogen, Carlsbad, USA). Lysates were incubated with α-peIF4E or α-m2A antibodies for 45 minutes. α-MNK1 antibody and IP buffer served as nonreactive and beads controls, respectively. Protein A Sepharose® slurry (Abcam, USA) was added (40 μL, 5 ng/μL) to each reaction and incubated overnight at 4°C on a rotary platform.

Following five washes with IP buffer, cross-links were reversed using Proteinase K (20 mg/ml) at 56°C for 40 minutes. After centrifugation at 12000 g for 1 min, RNA was isolated from supernatants, reverse transcribed, and BPXV M gene expression was quantified by qRT-PCR. Values were normalized to input RNA. All experiments were performed in triplicate.

### siRNA knockdown

Vero cells at ∼75% confluency were transfected with MAT2A-specific or negative control siRNA using Lipofectamine 3000 (Invitrogen, Carlsbad, USA) according to the manufacturer’s protocol. Target knockdown was confirmed by western blot analysis. Cell viability of siRNA-transfected cells was assessed by MTT assay. To evaluate the impact of gene knockdown on viral replication, transfected cells were infected with BPXV (MOI 5) at 48 hours post-transfection. After 1 hour of infection, cells were washed with PBS and incubated at 37°C. Infectious virus particles in culture supernatants were quantified by plaque assay at 48 hours post-infection. All experiments were performed in triplicate.

### Selection of DZNep-resistant virus variant

Vero cells were infected with BPXV (MOI 0.1) and maintained in DMEM containing either DZNep (1 µM) or DMSO (0.05%, vehicle control). Culture supernatants were harvested at 48-72 hpi (passage 1) and viral titers were determined by plaque assay. Sequential passages were performed by infecting fresh Vero cells with virus from the preceding passage under continuous drug pressure or vehicle control. Likewise, 50 such passages were carried out and the resulting virus(es) were termed as drBPXV (drug-resistant BPXV) and dsBPXV (drug-susceptible BPXV), respectively. To evaluate drug susceptibility, Vero cells were infected (MOI 0.1) with wild-type BPXV (P0; wtBPXV), drBPXV, or dsBPXV. After infection, cells were treated with DZNep (5 µM) or DMSO (0.05%). Viral titers in culture supernatants were determined by plaque assay at 48 hpi. All experiments were performed in triplicate.

### mRNA stability

Vero cells, in triplicates, were infected with BPXV at MOI of 5, followed by washing five times with PBS and addition of fresh DMEM. At 12 hpi, cells were treated with either 0.05% DMSO or 5 µM DZNep in the presence of 5 µg/ml Actinomycin D. At 0h, 1h and 2h post-drug treatment, the cells were scrapped and total RNA was isolated. The levels of BPXV *M* gene was quantified by qRT-PCR. The half-lives of the targeted mRNA was calculated as described previously^71^.

### RNA extraction, library preparation, and Illumina sequencing

Vero cells (80-85% confluent) were infected with drBPXV or dsBPXV (MOI 5), treated with DZNep (5 µM) at 1 hour post-infection, and harvested at 12 hpi for total RNA extraction using TRIzol reagent (Takara, China).

Poly(A) mRNA was enriched from total RNA (500 ng) using the NEBNext® Poly(A) mRNA Magnetic Isolation Module (New England Biolabs, MA, USA) according to the manufacturer’s protocol. Libraries were prepared using the NEBNext® Ultra™ II RNA Library Prep Kit. Enriched mRNA was fragmented in magnesium-based buffer at 94°C for 10 minutes using NEBNext® Random Primers to generate ∼300 nucleotide inserts. The fragmented RNA was reverse transcribed into first-strand cDNA and was followed by double-stranded DNA conversion and purification using 1.8X AMPure XP beads (Beckman Coulter, CA, USA). The purified dsDNA was subjected to end repair, 3’ adenylation, and loop adapter ligation.

Following USER enzyme treatment, adapter-ligated products were size-selected using AMPure XP beads to obtain 400-600 bp libraries. Libraries were amplified through 12 PCR cycles using NEBNext® Ultra II Q5 Master Mix and Multiplex Oligos, then purified with 0.9X AMPure XP beads and eluted in 15 µl of 0.1X TE buffer.

Adapter ligated products were treated with USER enzyme and size-selected using AMPure XP beads (Beckman Coulter, CA, USA) to target a library size of 400–600 bp. The cDNA libraries were amplified through 12 PCR cycles using NEBNext® Ultra II Q5 Master Mix and NEBNext® Multiplex Oligos for Illumina. The amplified libraries were purified with 0.9X AMPure XP beads and eluted in 15 µl of 0.1X TE buffer.

Library quality was assessed using a Qubit Fluorometer (Invitrogen, Life Technologies, USA) for quantification and Tapestation system (Agilent Technologies, USA) with HSDNA kit for size distribution analysis. Pooled libraries underwent cluster generation on the c-Bot system followed by paired-end sequencing (2×150 bp) on an Illumina HiSeq X10. Demultiplexing and adapter trimming were performed using CASAVA v1.8.2 (Illumina Inc.).

### RNA-seq data analysis

Raw read quality was assessed using FastQC (v0.11.8) to examine base quality score distribution, sequence quality score distribution, average base content per read, and GC content distribution. Adapter sequences (AGATCGGAAGAGC) were removed using Trim Galore (v0.6.2), which also performed automated quality trimming. Reads shorter than 20 bp or with low-quality ends (Phred score <20) were discarded. The *Chlorocebus sabaeus* reference genome (Ensembl release 110) and BPXV genome (NCBI accession no. MW883892.1) were indexed using BWA (v0.7.17), and preprocessed reads were aligned using BWA-MEM with default parameters. Mapped reads were quantified at the gene level using Samtools (v0.1.19).

Differential expression analysis was performed using DESeq (v1) to identify genes with significant expression changes. Differentially expressed genes (DEGs) were defined using threshold criteria of fold change ≥2 and p-value <0.05. DEGs were functionally annotated using BLASTx (v2.2.29+) against the NR database, with annotations retrieved from UniProt and KEGG databases. Gene Ontology (GO) terms were visualized using WEGO with a log10-scaled y-axis. Expression patterns were visualized through volcano plots and MA plots generated using custom R scripts, while heatmaps were created using MeV software (v4.8.1).

### Homology modelling and molecular dynamics simulation

Homology modelling was performed to resolve missing residues (142KRGGNE147) in the crystal structures of VP39 bound with SAH (PDB id 1VP3) and SAM (PDB id 1VPT). The top 5 modelled structures were aligned with respective crystal structures to obtain accurate ligand coordinates. Missing residues and hydrogens were added, and the proteins were parameterized using the AMBER force field. Ligand topology and parameters were generated using CGenFF and ACPYPE^72^.

The protein-ligand complexes were placed in cubic TIP3PBOX (85.266 × 85.285 × 94.169 Å) and neutralized with appropriate ions. Energy minimization was performed using the steepest descent method until the maximum force was below 1000 kJ/mol/nm. System equilibration was conducted in two stages: NVT ensemble (200 ps, 300 K) using the Berendsen thermostat, followed by NPT ensemble (200 ps, 1 bar) using the Parrinello-Rahman barostat. Position restraints were applied to the protein and ligand during equilibration.

Unrestrained MD simulations were conducted for 200 ns in the NPT ensemble (300 K, 1 bar) with a 2 fs integration time step and periodic boundary conditions. Long-range electrostatic interactions were calculated using the Particle Mesh Ewald method (cut-off: 1.70 nm), and van der Waals interactions had a cut-off distance of 1.7 nm. Coordinates and velocities were recorded every 100 ps for trajectory analysis.

Trajectory analysis was performed using MD Analysis and VMD, focusing on root mean square deviation (RMSD), root mean square fluctuation (RMSF), and hydrogen bonding interactions to assess the stability and conformational behaviour of the protein-ligand complexes.

### Direct competition assay for viral fitness

Direct competition assay was conducted as previously described^73^. drBPXV and dsBPXV variants were mixed in ratios of 9:1, 1:1, and 1:9, and used to infect Vero cells at an MOI of 0.1. At 1 hour post-infection, cells were washed with PBS and supplemented with fresh DMEM. Upon observing cytopathic effects, supernatants were harvested and passaged sequentially three additional times. At each passage, viral DNA was isolated from the supernatant using a Qiagen DNeasy column (Qiagen, Hilden, Germany). The VP39 region was amplified by PCR, and the purified PCR product was subjected to Sanger sequencing. Resulting chromatograms for each group were analyzed using Geneious Prime software (v 2023.2.1).

### Cloning and overexpression of VP39

Wild-type and mutant (M236I) BPXV VP39 genes were cloned into pDONR221 vector and subsequently transferred to pDEST40 vector using GATEWAY cloning technology (Invitrogen, Carlsbad, CA, USA) according to manufacturer’s instructions. Constructs were verified by sequencing. HeLa cells (for BPXV, VACV, and LSDV) or A549 cells (for SARS-CoV-2 delta variant) in 6-well plates were transfected with 2 μg of pDEST40:VP39_wt or pDEST40:VP39_mut using Lipofectamine 3000. At 48 hours post-transfection, cells were infected with respective viruses at an MOI of 5. After PBS washing, cells were incubated at 37°C. Infectious virus particles in supernatants were quantified at 48 hours post-infection as described in section 3.1.

### Expression and purification of VP39

VP39_wt and VP39_mut were subcloned from pDONR221 into pDEST17 vectors using GATEWAY cloning technology (Invitrogen, Carlsbad, CA, USA). BL21-A1 cells transformed with these constructs were induced with 0.2% L-arabinose for 3 hours. Cell pellets were resuspended in Xtractor lysis buffer (TaKaRa Bio, Ohtsu, Japan) supplemented with Benzonase nuclease, lysozyme, and EDTA-free protease inhibitor cocktail. Cells were lysed by sonication (40% power, 10-second cycles, 60 seconds total), and debris was removed by centrifugation (12,000 g, 30 minutes).

Clarified lysates were incubated with Ni-NTA agarose beads and loaded onto polypropylene columns (Qiagen). Non-specific proteins were removed by washing with His-select® washing buffer, and target proteins were eluted using His-select® elution buffer (Sigma-Aldrich). Purified proteins were analyzed by 8% SDS-PAGE and Western blot using anti-His antibody (Invitrogen). Proteins were concentrated and buffer-exchanged using Amicon® Ultra Centrifugal Filters (30 kDa MWCO) and stored at-20°C in storage buffer (20 mM Tris-HCl, 100 mM NaCl, 1 mM DTT, 0.1 mM EDTA, 0.1% Triton X-100, 50% glycerol, pH 8.0). Protein concentrations were determined using the Qubit Protein Assay Kit (Invitrogen). All purification steps were performed at 4°C.

### *In vitro* transcription and mRNA capping

IL6-Nanoluciferase was amplified from plasmid #84394 (Addgene) using primers containing T7 promoter and Kozak sequences (forward) and T30 sequence (reverse). Amplicons were purified using the Monarch PCR Cleanup Kit (New England Biolabs). mRNA was synthesized by *in vitro* transcription using the HiScribe T7 High Yield RNA Synthesis Kit (New England Biolabs) according to manufacturer’s instructions, except for the replacement of 100% UTP with N1-Methylpseudouridine-5’-Triphosphate (N1MePsU, TriLink BioTechnologies).

The transcribed RNA was DNase I-treated and purified using the MEGAclear Transcription Clean-Up Kit (Thermo Fisher Scientific). RNA yield was quantified using the Qubit RNA HS Assay Kit (Invitrogen). Cap-0 modification of mRNA constructs was performed using vaccinia capping enzyme (New England Biolabs) according to manufacturer’s instruction. For cap-1 modification, 800 ng of purified VP39_wt or VP39_mut MTase (normalized by Qubit protein assay kit, Invitrogen) was added in the capping reaction and incubated for 1 hour at 37℃. Modified mRNA was purified using MEGAclear Transcription Clean-Up Kit and quantified using Qubit RNA HS Assay Kit (Invitrogen).

### Time-course IL6-Nanoluciferase secretion

HeLa cells were cultured in FluoroBrite DMEM (Invitrogen) supplemented with 10% FBS and L-glutamine in 96-well plate. At 70 - 80% confluence, cells were transfected in triplicates with 200 ng of cap-0, cap-0 + VP39_wt-capped, or cap-0 + VP39_mut-capped mRNA using Lipofectamine MessengerMAX (Invitrogen) according to the manufacturer’s protocol. At 6 hours post-transfection, transfection media was replaced with fresh FluoroBrite DMEM containing L-glutamine and 10% FBS. Each day, complete 150 µl of old media were harvested from each well and replaced with fresh media supplemented with 10% FBS and L-glutamine. For luminescence activity 10 µl of harvested media was diluted in Nano-Glo luciferase assay system (Promega) according to manufacturer’s instructions. Readings were measured in a black-walled clear bottom 96 well plate (Costar) to avoid crosstalk using Spectramax i3x (Molecular Devices) in end-point mode.

### Polysome profiling

Polysome profiling was performed as previously described with some modifications^74^. Vero cells were infected with drBPXV or dsBPXV (MOI 5) for 12 hours, with cycloheximide (100 µg/ml) added for the final 10 minutes. Cells were lysed in polysome lysis buffer (5 mM Tris-HCl pH 7.5, 2.5 mM MgCl2, 1.5 mM KCl, 100 µg/ml cycloheximide, 1 mM DTT, 1% Triton X, 40 U/ml RNase inhibitor, 1x EDTA-free protease inhibitor cocktail) on ice for 30 min followed by centrifugation at 12,000g for 20 min. The resulting supernatant was collected and stored at-80°C till use.

Lysates (5 OD units) were layered on 15-40% sucrose gradients (20 mM Tris-HCl pH 7.5, 10 mM MgCl₂, 50 mM KCl, 100 mg/ml cycloheximide, 1 mM DTT, 20 U/ml RNase inhibitor) and centrifuged at 39,000 rpm for 2 hours at 4°C (Beckman Ultracentrifuge, SW41 rotor). Gradients were fractionated (0.6 ml portions) using a Piston Gradient Fractionator (Biocomp Instruments). RNA was extracted from 0.2 ml of each fraction (starting from 80S ribosomal subunit) using TRIzol. cDNA libraries were prepared using oligo(dT) primers, and BPXV mRNA and β-actin expression were quantified by qRT-PCR. Percent distribution of BPXV mRNA across the gradient was determined using the ΔΔCt method^74^.

