## Supplementary figures for "DZNep-induced single point mutation (M236I) in poxviral 2’-O-methyltransferase enhances mRNA stability and translation efficiency"

### Slide 1
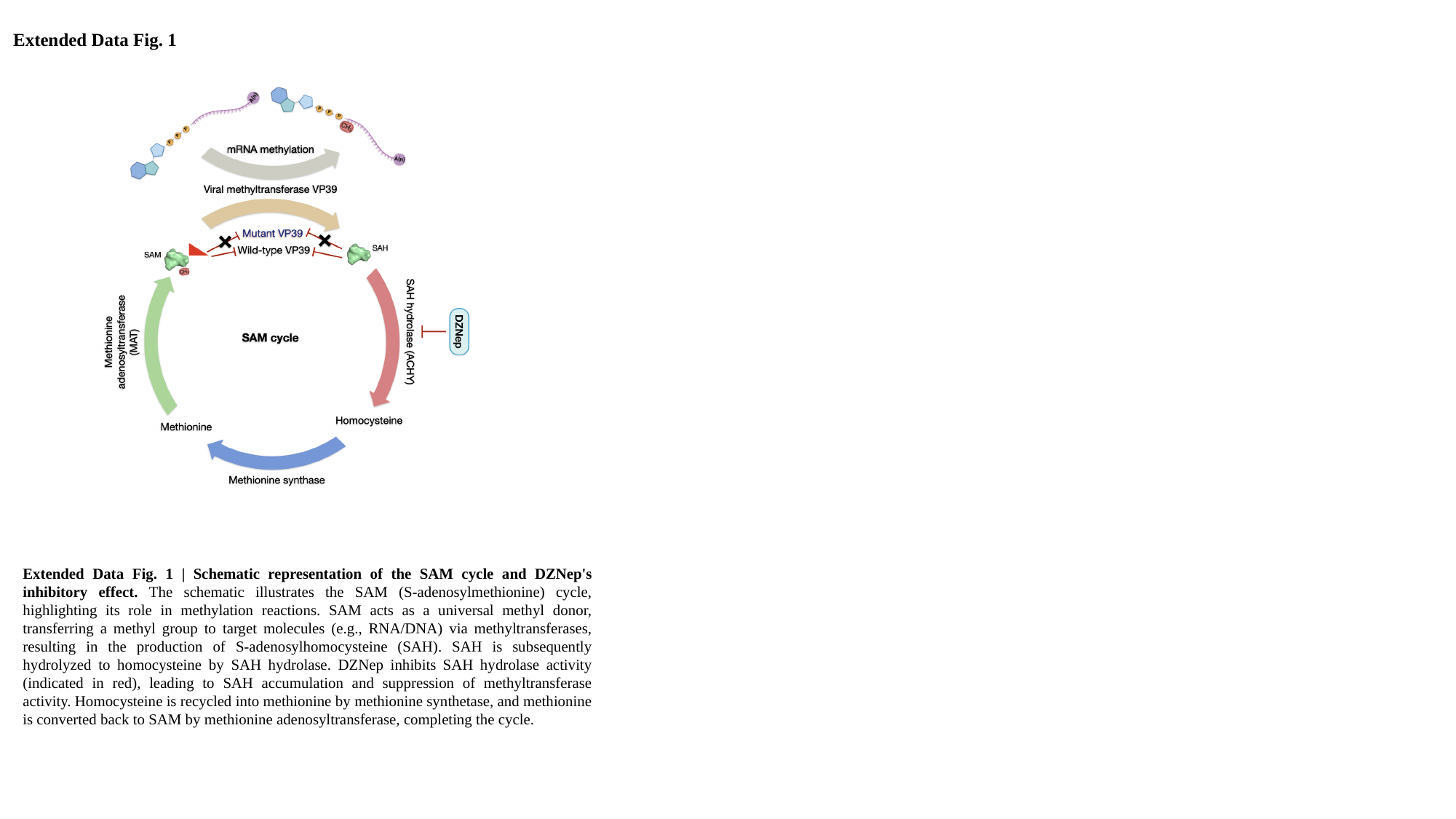

Extended Data Fig. 1
Extended Data Fig. 1 | Schematic representation of the SAM cycle and DZNep's inhibitory effect. The schematic illustrates the SAM (S-adenosylmethionine) cycle, highlighting its role in methylation reactions. SAM acts as a universal methyl donor, transferring a methyl group to target molecules (e.g., RNA/DNA) via methyltransferases, resulting in the production of S-adenosylhomocysteine (SAH). SAH is subsequently hydrolyzed to homocysteine by SAH hydrolase. DZNep inhibits SAH hydrolase activity (indicated in red), leading to SAH accumulation and suppression of methyltransferase activity. Homocysteine is recycled into methionine by methionine synthetase, and methionine is converted back to SAM by methionine adenosyltransferase, completing the cycle.

### Slide 2
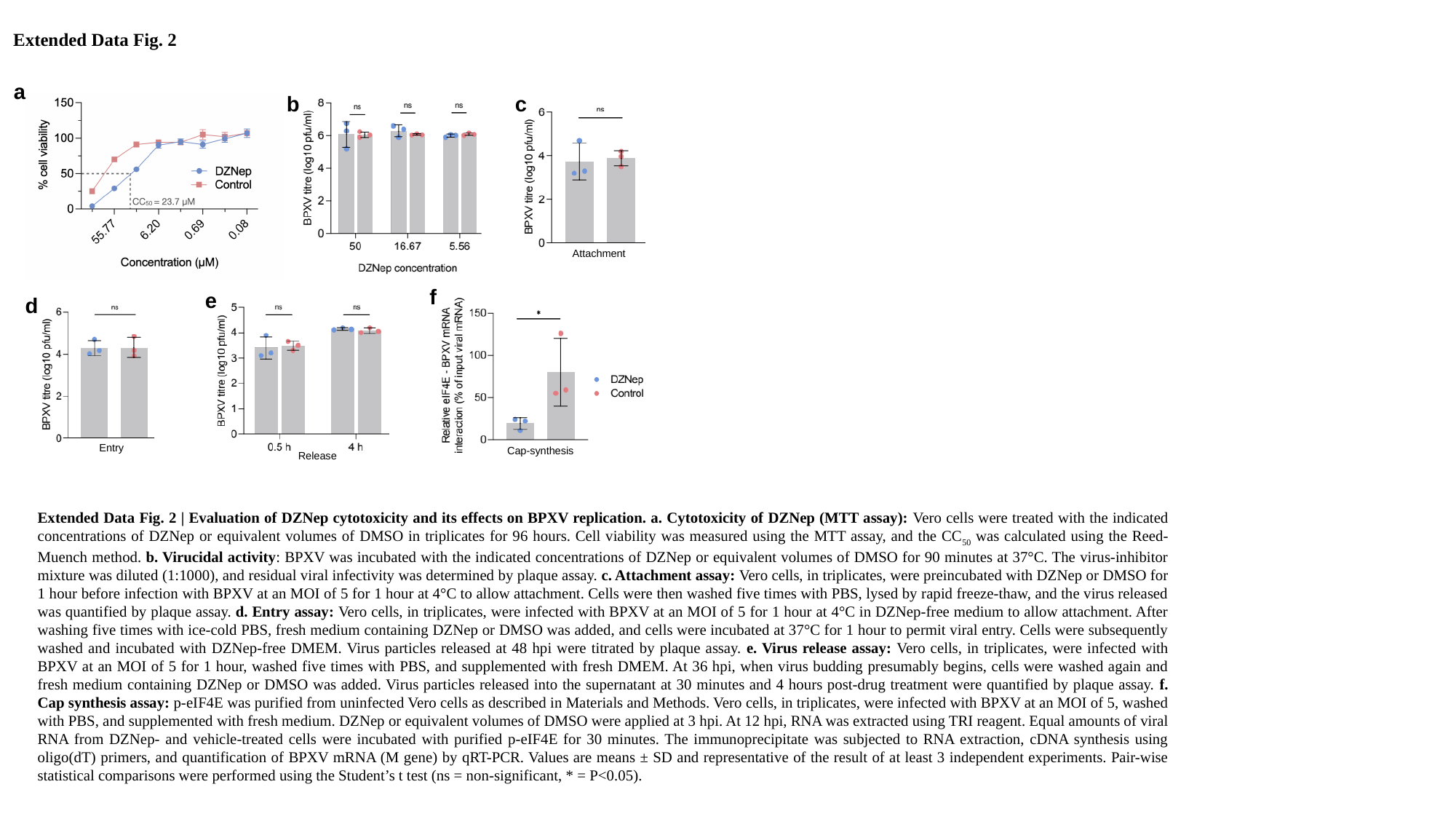

Extended Data Fig. 2
a
b
c
Attachment
f
Cap-synthesis
e
Release
d
Entry
Extended Data Fig. 2 | Evaluation of DZNep cytotoxicity and its effects on BPXV replication. a. Cytotoxicity of DZNep (MTT assay): Vero cells were treated with the indicated concentrations of DZNep or equivalent volumes of DMSO in triplicates for 96 hours. Cell viability was measured using the MTT assay, and the CC_50_ was calculated using the Reed-Muench method. b. Virucidal activity: BPXV was incubated with the indicated concentrations of DZNep or equivalent volumes of DMSO for 90 minutes at 37°C. The virus-inhibitor mixture was diluted (1:1000), and residual viral infectivity was determined by plaque assay. c. Attachment assay: Vero cells, in triplicates, were preincubated with DZNep or DMSO for 1 hour before infection with BPXV at an MOI of 5 for 1 hour at 4°C to allow attachment. Cells were then washed five times with PBS, lysed by rapid freeze-thaw, and the virus released was quantified by plaque assay. d. Entry assay: Vero cells, in triplicates, were infected with BPXV at an MOI of 5 for 1 hour at 4°C in DZNep-free medium to allow attachment. After washing five times with ice-cold PBS, fresh medium containing DZNep or DMSO was added, and cells were incubated at 37°C for 1 hour to permit viral entry. Cells were subsequently washed and incubated with DZNep-free DMEM. Virus particles released at 48 hpi were titrated by plaque assay. e. Virus release assay: Vero cells, in triplicates, were infected with BPXV at an MOI of 5 for 1 hour, washed five times with PBS, and supplemented with fresh DMEM. At 36 hpi, when virus budding presumably begins, cells were washed again and fresh medium containing DZNep or DMSO was added. Virus particles released into the supernatant at 30 minutes and 4 hours post-drug treatment were quantified by plaque assay. f. Cap synthesis assay: p-eIF4E was purified from uninfected Vero cells as described in Materials and Methods. Vero cells, in triplicates, were infected with BPXV at an MOI of 5, washed with PBS, and supplemented with fresh medium. DZNep or equivalent volumes of DMSO were applied at 3 hpi. At 12 hpi, RNA was extracted using TRI reagent. Equal amounts of viral RNA from DZNep- and vehicle-treated cells were incubated with purified p-eIF4E for 30 minutes. The immunoprecipitate was subjected to RNA extraction, cDNA synthesis using oligo(dT) primers, and quantification of BPXV mRNA (M gene) by qRT-PCR. Values are means ± SD and representative of the result of at least 3 independent experiments. Pair-wise statistical comparisons were performed using the Student’s t test (ns = non-significant, * = P<0.05).

### Slide 3
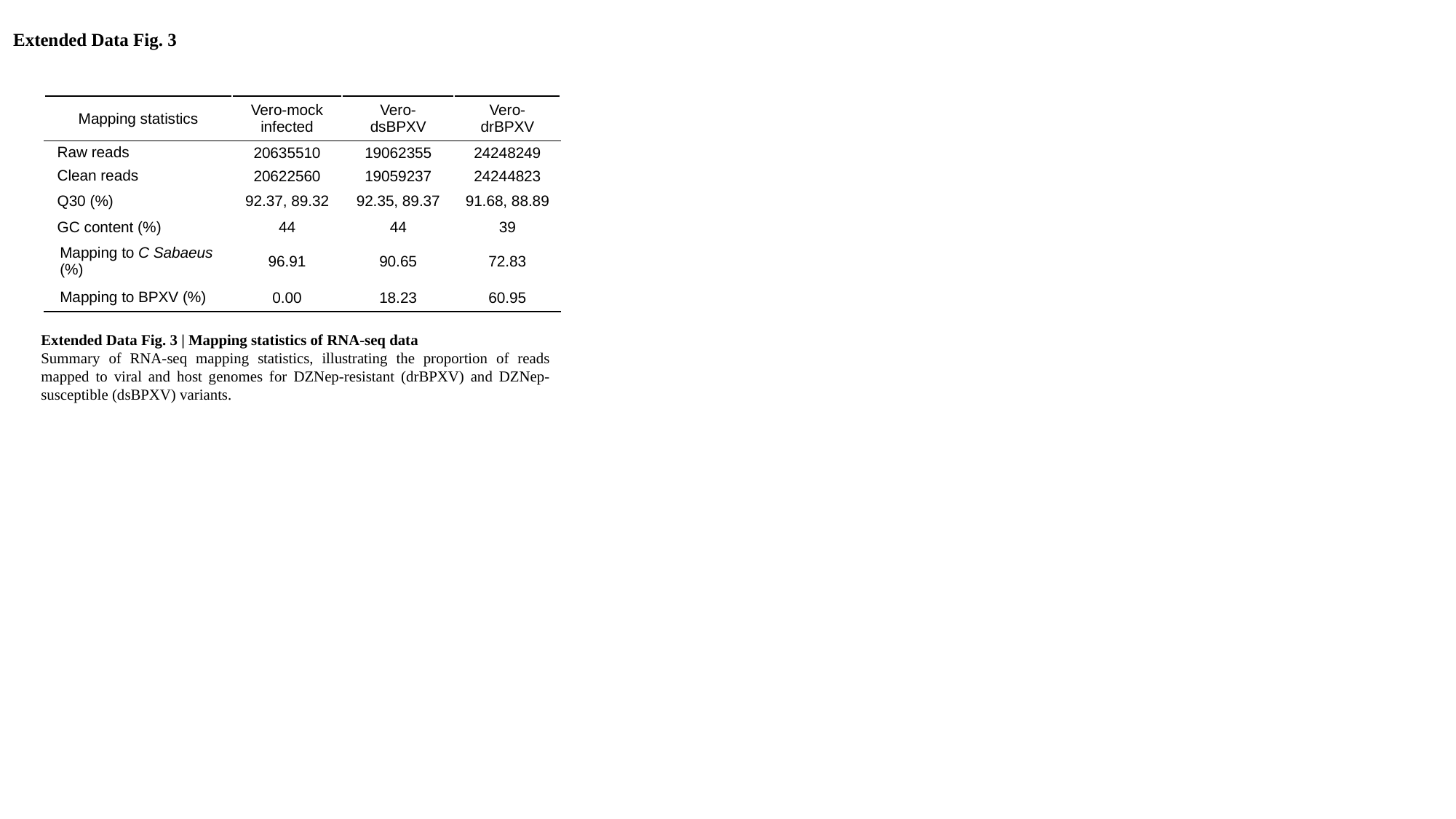

Extended Data Fig. 3
| Mapping statistics | Vero-mock infected | Vero-dsBPXV | Vero-drBPXV |
| --- | --- | --- | --- |
| Raw reads | 20635510 | 19062355 | 24248249 |
| Clean reads | 20622560 | 19059237 | 24244823 |
| Q30 (%) | 92.37, 89.32 | 92.35, 89.37 | 91.68, 88.89 |
| GC content (%) | 44 | 44 | 39 |
| Mapping to C Sabaeus (%) | 96.91 | 90.65 | 72.83 |
| Mapping to BPXV (%) | 0.00 | 18.23 | 60.95 |
Extended Data Fig. 3 | Mapping statistics of RNA-seq data
Summary of RNA-seq mapping statistics, illustrating the proportion of reads mapped to viral and host genomes for DZNep-resistant (drBPXV) and DZNep-susceptible (dsBPXV) variants.

### Slide 4
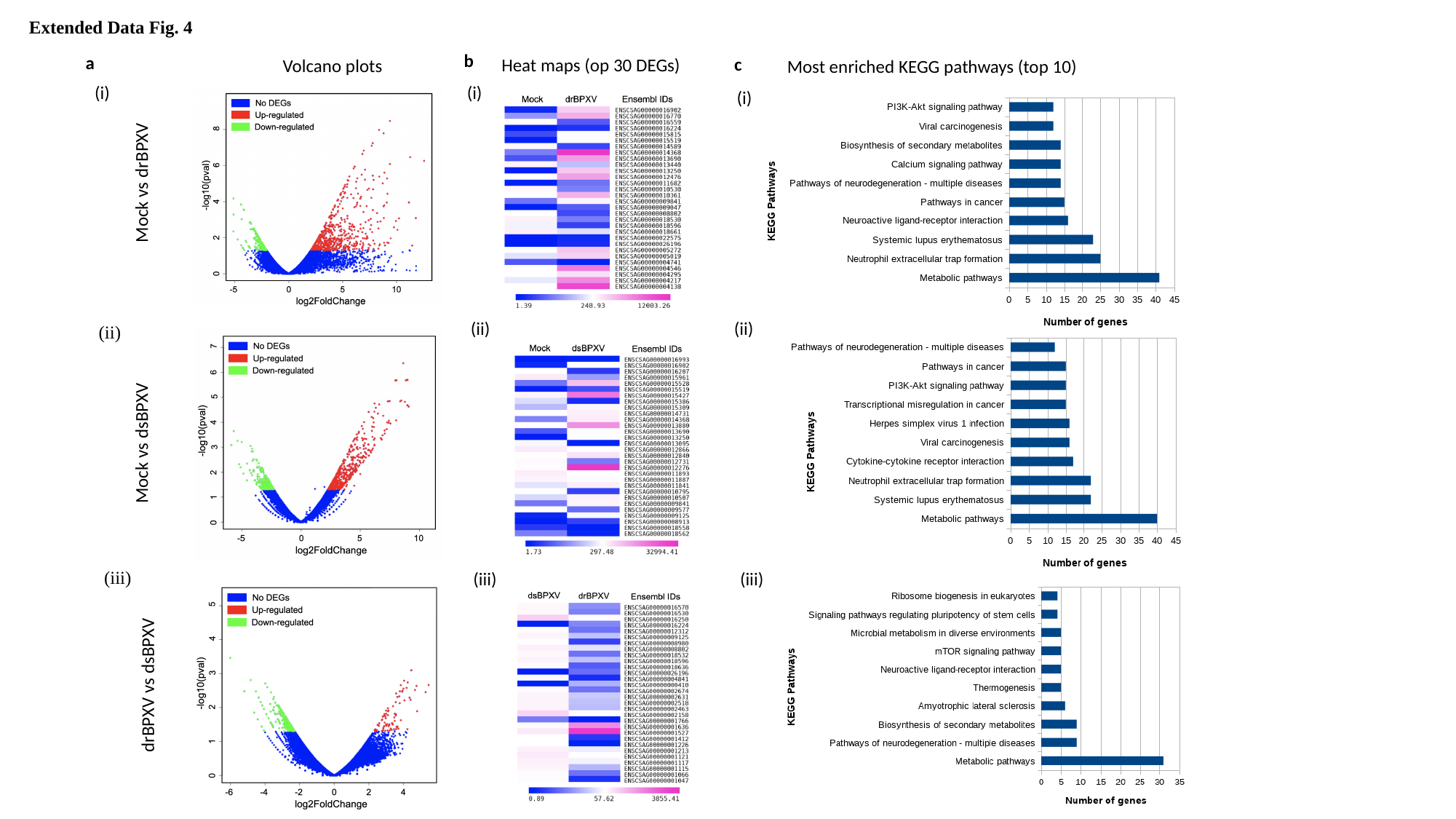

Extended Data Fig. 4
b
Heat maps (op 30 DEGs)
(i)
(ii)
(iii)
a
Volcano plots
(i)
Mock vs drBPXV
(ii)
Mock vs dsBPXV
(iii)
drBPXV vs dsBPXV
c
Most enriched KEGG pathways (top 10)
(i)
(ii)
(iii)

### Slide 5
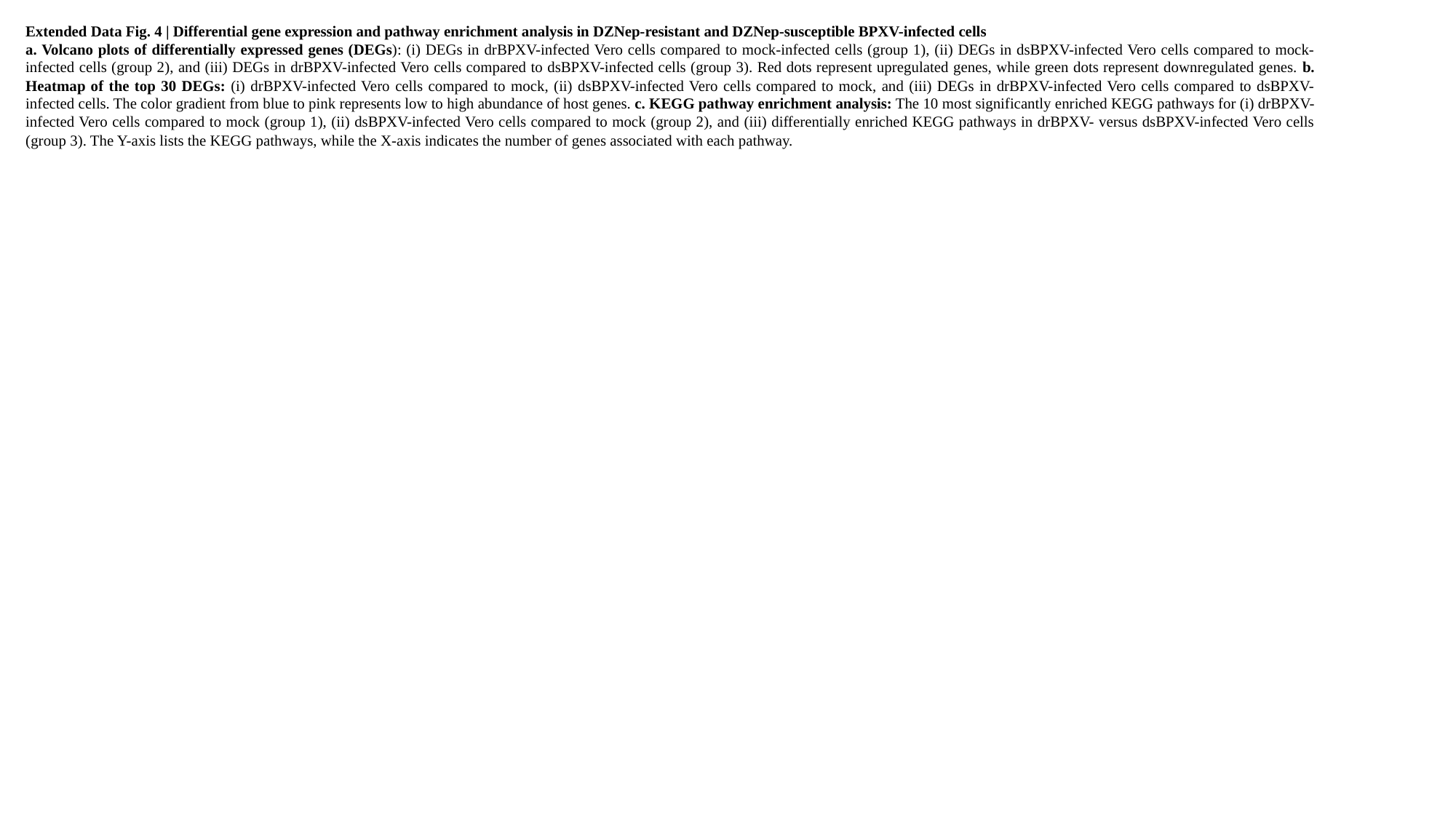

Extended Data Fig. 4 | Differential gene expression and pathway enrichment analysis in DZNep-resistant and DZNep-susceptible BPXV-infected cells
a. Volcano plots of differentially expressed genes (DEGs): (i) DEGs in drBPXV-infected Vero cells compared to mock-infected cells (group 1), (ii) DEGs in dsBPXV-infected Vero cells compared to mock-infected cells (group 2), and (iii) DEGs in drBPXV-infected Vero cells compared to dsBPXV-infected cells (group 3). Red dots represent upregulated genes, while green dots represent downregulated genes. b. Heatmap of the top 30 DEGs: (i) drBPXV-infected Vero cells compared to mock, (ii) dsBPXV-infected Vero cells compared to mock, and (iii) DEGs in drBPXV-infected Vero cells compared to dsBPXV-infected cells. The color gradient from blue to pink represents low to high abundance of host genes. c. KEGG pathway enrichment analysis: The 10 most significantly enriched KEGG pathways for (i) drBPXV-infected Vero cells compared to mock (group 1), (ii) dsBPXV-infected Vero cells compared to mock (group 2), and (iii) differentially enriched KEGG pathways in drBPXV- versus dsBPXV-infected Vero cells (group 3). The Y-axis lists the KEGG pathways, while the X-axis indicates the number of genes associated with each pathway.

### Slide 6
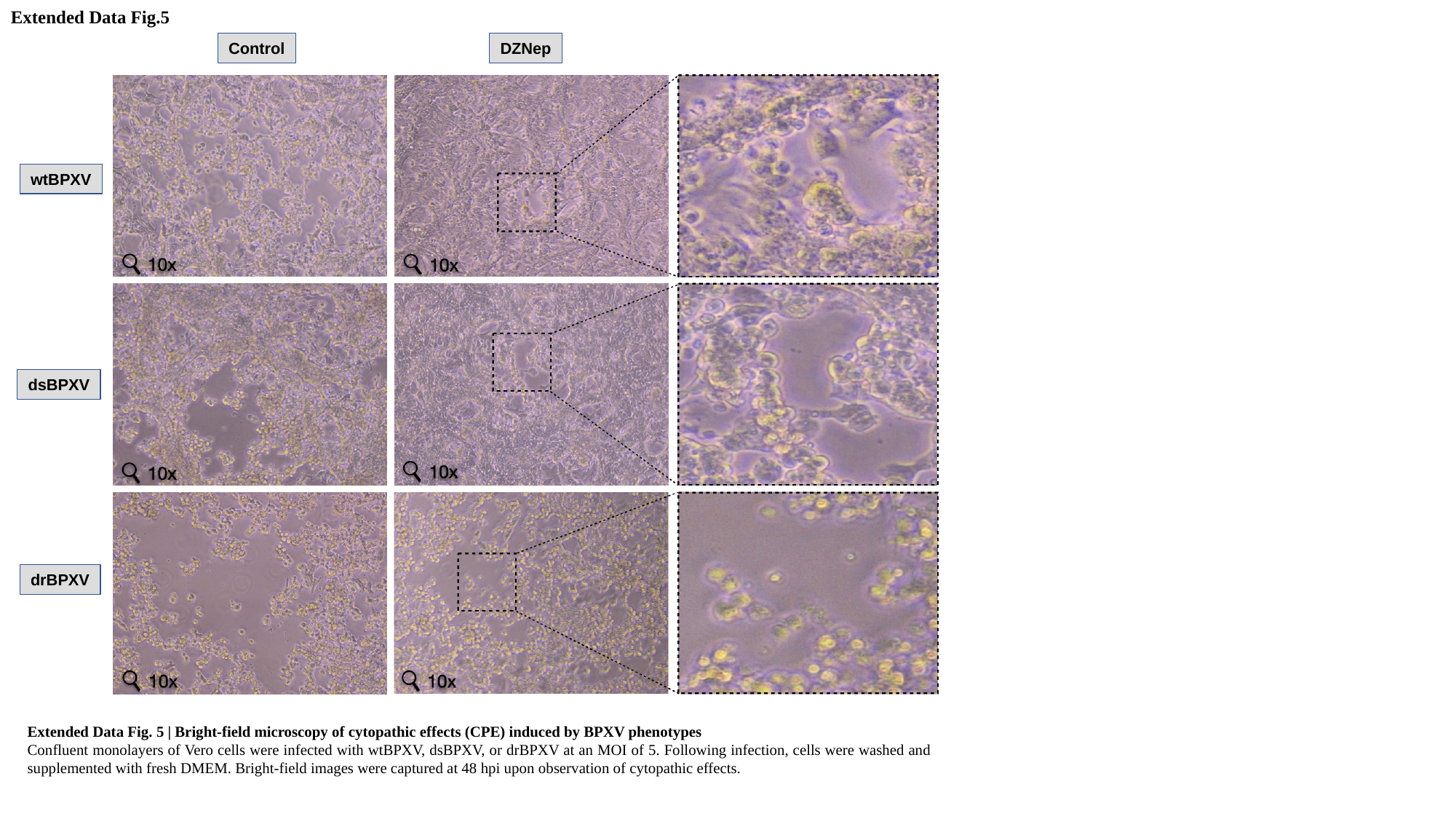

Extended Data Fig.5
Control
DZNep
wtBPXV
dsBPXV
drBPXV
Extended Data Fig. 5 | Bright-field microscopy of cytopathic effects (CPE) induced by BPXV phenotypes
Confluent monolayers of Vero cells were infected with wtBPXV, dsBPXV, or drBPXV at an MOI of 5. Following infection, cells were washed and supplemented with fresh DMEM. Bright-field images were captured at 48 hpi upon observation of cytopathic effects.

### Slide 7
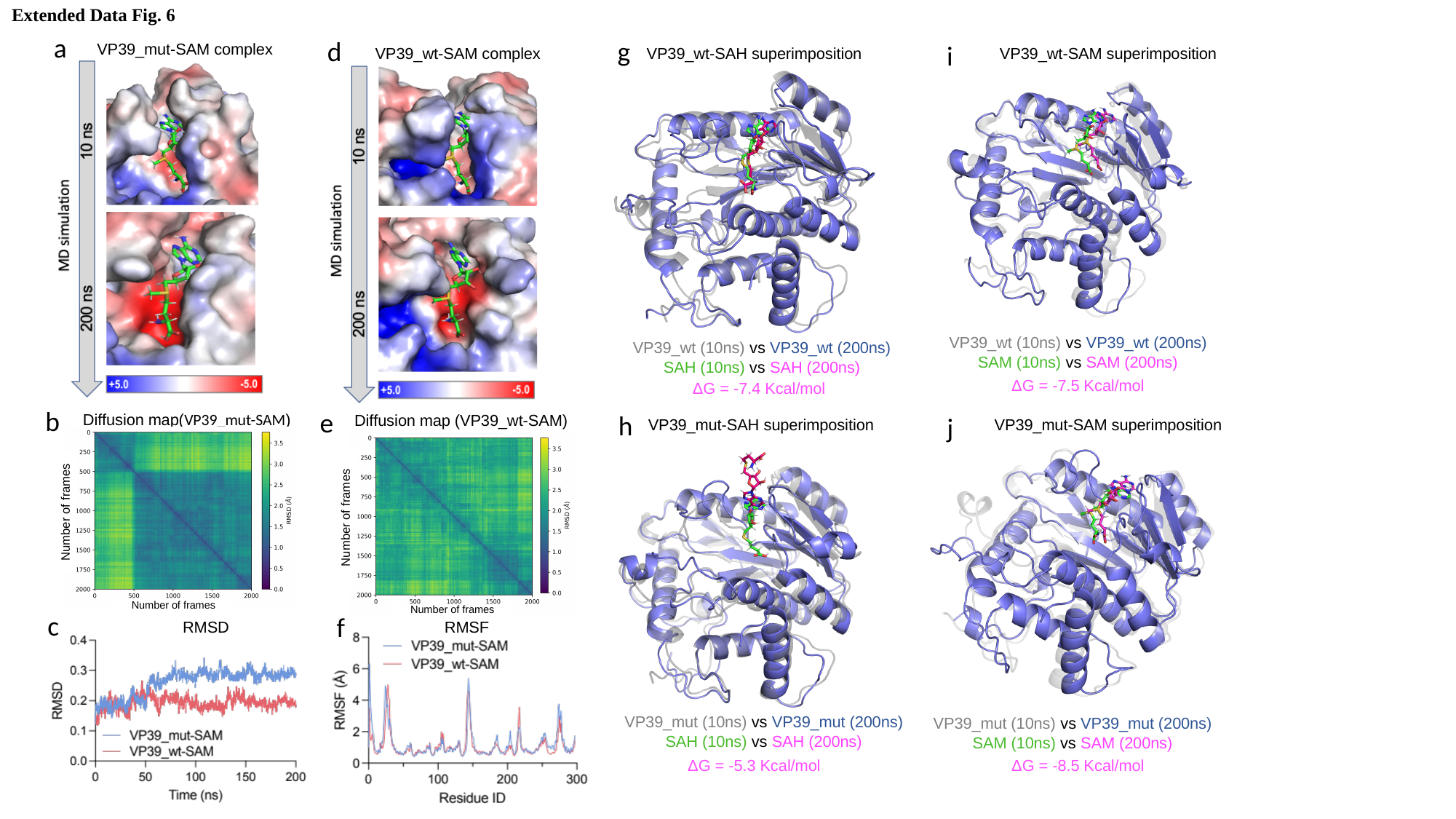

Extended Data Fig. 6
a
VP39_mut-SAM complex
b
Diffusion map(VP39_mut-SAM)
Number of frames
Number of frames
d
VP39_wt-SAM complex
Diffusion map (VP39_wt-SAM)
e
Number of frames
Number of frames
g
VP39_wt-SAH superimposition
VP39_wt (10ns) vs VP39_wt (200ns)
SAH (10ns) vs SAH (200ns)
ΔG = -7.4 Kcal/mol
i
VP39_wt-SAM superimposition
VP39_wt (10ns) vs VP39_wt (200ns)
SAM (10ns) vs SAM (200ns)
ΔG = -7.5 Kcal/mol
h
VP39_mut-SAH superimposition
VP39_mut (10ns) vs VP39_mut (200ns)
SAH (10ns) vs SAH (200ns)
j
VP39_mut-SAM superimposition
VP39_mut (10ns) vs VP39_mut (200ns)
SAM (10ns) vs SAM (200ns)
ΔG = -8.5 Kcal/mol
c
RMSD
f
RMSF
ΔG = -5.3 Kcal/mol

### Slide 8
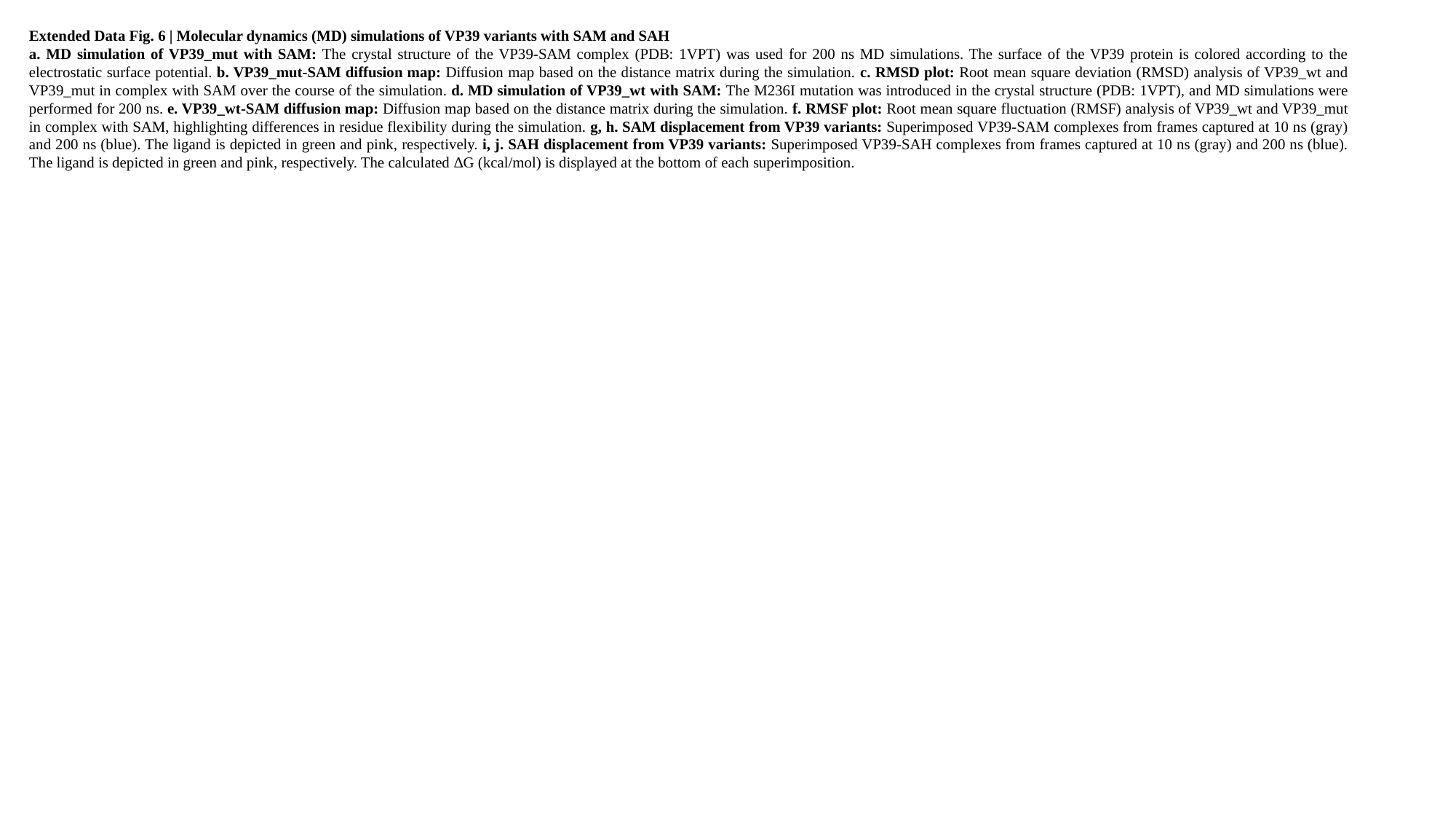

Extended Data Fig. 6 | Molecular dynamics (MD) simulations of VP39 variants with SAM and SAH
a. MD simulation of VP39_mut with SAM: The crystal structure of the VP39-SAM complex (PDB: 1VPT) was used for 200 ns MD simulations. The surface of the VP39 protein is colored according to the electrostatic surface potential. b. VP39_mut-SAM diffusion map: Diffusion map based on the distance matrix during the simulation. c. RMSD plot: Root mean square deviation (RMSD) analysis of VP39_wt and VP39_mut in complex with SAM over the course of the simulation. d. MD simulation of VP39_wt with SAM: The M236I mutation was introduced in the crystal structure (PDB: 1VPT), and MD simulations were performed for 200 ns. e. VP39_wt-SAM diffusion map: Diffusion map based on the distance matrix during the simulation. f. RMSF plot: Root mean square fluctuation (RMSF) analysis of VP39_wt and VP39_mut in complex with SAM, highlighting differences in residue flexibility during the simulation. g, h. SAM displacement from VP39 variants: Superimposed VP39-SAM complexes from frames captured at 10 ns (gray) and 200 ns (blue). The ligand is depicted in green and pink, respectively. i, j. SAH displacement from VP39 variants: Superimposed VP39-SAH complexes from frames captured at 10 ns (gray) and 200 ns (blue). The ligand is depicted in green and pink, respectively. The calculated ΔG (kcal/mol) is displayed at the bottom of each superimposition.

### Slide 9
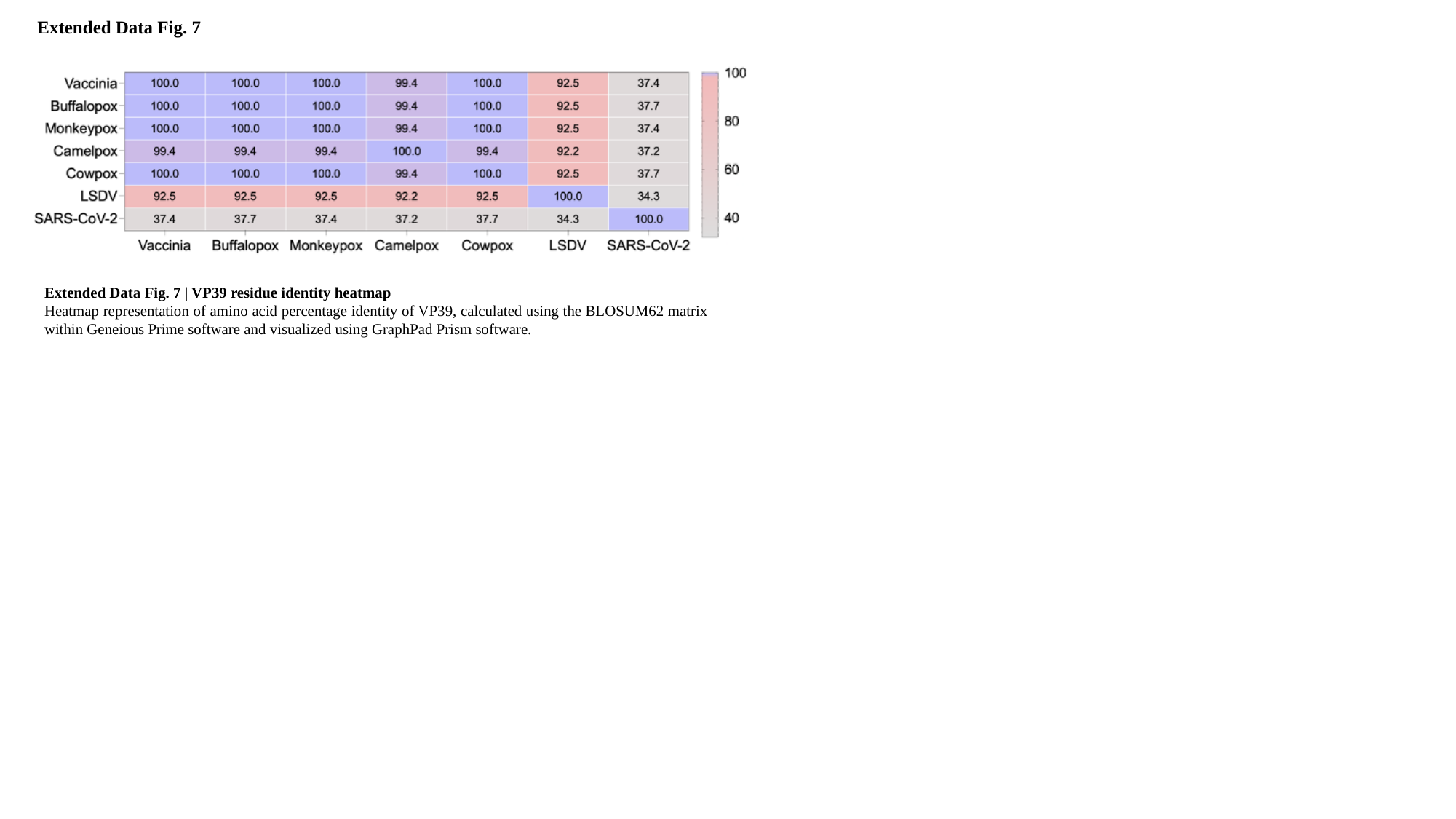

Extended Data Fig. 7
Extended Data Fig. 7 | VP39 residue identity heatmap
Heatmap representation of amino acid percentage identity of VP39, calculated using the BLOSUM62 matrix within Geneious Prime software and visualized using GraphPad Prism software.
